# Spatio-temporal transcriptomic analysis reveals distinct nephrotoxicity, DNA damage and regeneration response after cisplatin

**DOI:** 10.1101/2023.01.03.522568

**Authors:** Lukas S. Wijaya, Steven J. Kunnen, Panuwat Trairatphisan, Ciaran Fisher, Meredith E. Crosby, Kai Schaefer, Karen Bodie, Erin E. Vaughan, Laura Breidenbach, Thomas Reich, Diana Clausznitzer, Sylvestre A. Bonnet, Sipeng Zheng, Chantal Pont, James L. Stevens, Sylvia Le Dévédec, Bob van de Water

## Abstract

Nephrotoxicity caused by drug or chemical exposure involves different mechanisms and nephron segments as well as a complex temporal integration of injury and repair responses. Distinct cellular transcriptional programs regulate the time-dependent tissue injury and regeneration responses. Whole kidney transcriptome analysis cannot dissect the complex the nephron segment spatio- temporal injury and regeneration responses. Here, we used laser capture microdissection of formalin-fixed paraffin embedded sections followed by whole genome targeted RNA-sequencing-TempO-Seq and co-expression gene-network (module) analysis to determine the spatial-temporal responses in rat kidney glomeruli (GM), cortical proximal tubules (CPT) and outer-medulla proximal tubules (OMPT) comparison with whole kidney, after a single dose of the nephrotoxicant cisplatin. We demonstrate that cisplatin induced early onset of DNA damage in both CPT and OMPT, but not GM. Sustained DNA damage response was strongest in OMPT coinciding with OMPT specific inflammatory signaling, actin cytoskeletal remodeling and increased glycolytic metabolism coincident with suppression of mitochondrial activity. Later responses reflected regeneration-related cell cycle pathway activation and ribosomal biogenesis in the injured OMPT regions. Activation of modules containing kidney injury biomarkers was strongest in the OMPT, with OMPT *Clu* expression best correlating with urinary clusterin biomarker measurements compared the correlation of Kim1. Our findings also showed that whole kidney responses were less sensitive than OMPT. In conclusion, our LCM-TempO-Seq method reveals a detailed spatial mechanistic understanding of renal injury/regeneration after nephrotoxicant exposure and identifies the most representative mechanism-based nephron segment specific renal injury biomarkers.

## Introduction

Drug-induced kidney injury (DIKI) remains a major problem for drug development and clinical safety management due to poor understanding of the underlying mechanisms^1–4^. Nephron segments differ in sensitivity to nephrotoxicants and involve specific mechanisms that underlie segment-specific responses^5–7^. Given its physiological role, the proximal tubule is a sensitive target for nephrotoxicants^8^. Toxicogenomics assessment of whole kidney responses has typically been applied to derive mechanistic insights in nephrotoxicity^9^. However, given the cellular heterogeneity of the kidney and the differential sensitivity of nephron segments tor renal toxicants, characterizing molecular mechanisms of DIKI onset and progression at the level of the entire kidney transcriptome does not provide insights into segment-specific responses. Rather, the whole kidney transcriptomic response represents an average of the gene expression responses across different nephron segments and does not reveal the complexity of segment-specific responses to injury progression and repair.

To overcome these limitations and advance the mechanistic understanding of spatio-temoral responses during DIKI we focused on the assessment of nephron segment specific responses using laser capture microdissection (LCM) in combination with targeted RNA-sequencing of formalin-fixed material based on the TempO-Seq technology^10^. We have applied this approach to understand the segment-specific transcriptional response of cisplatin. Cisplatin is a widely used chemotherapeutic agent to treat various cancers^11^. However, one third of the patients receiving cisplatin develop renal adverse effects^12^. The proximal tubule cells are the primary target in both pre-clinical species and human patients due to the high expression of the organic cationic transporter 2 (OCT2)^13–15^ and increased active transport of cisplatin into the proximal tubule epithelial cells^16,17^. The high intracellular cisplatin concentration in proximal tubule cells results in DNA damage in association with mitochondrial toxicity, cytoskeletal reorganization and inflammation which contribute to proximal tubular cell death resulting in acute renal failure^15^. Acute renal failure due to proximal tubular injury after cisplatin treatment is followed by tissue regeneration and degeneration; the balance between acute injury and regeneration responses governs the longer-term impact on overall renal function^15^. While these responses have been observed at the transcriptome level in the whole kidney the contribution of distinct mechanisms of injury and regeneration across different segments remains obscured. Finally, while cisplatin induces the expression of novel renal injury biomarkers related to proximal tubular injury such as KIM1, clusterin, and NGAL^18^, it is unclear whether these biomarkers reflect the responses of the (most) affected segments or are general kidney response to loss of function.

Herein we conducted a systematic spatial-temporal study in rat *in vivo* to map the nephron-segment specific cisplatin responses. We first established and validated the LCM-TempO-Seq protocol and mapped the transcriptional spatiotemporal dynamics in different nephron segments upon cisplatin exposure. To interpret the biological context of segment-specific transcriptomic data, we leveraged a recently established Kidney TXG-MAPr application (manuscript in preparation; https://txg-mapr.eu/). This platform was established by a data-driven weighted gene co-expression network analysis (WGCNA) based on a large TG-GATEs rat transcriptomics dataset from rat kidney after nephrotoxicant treatment with different doses and time points^9^. The DIKI-TXG-MAPr facilitates analysis of renal pathology-relevant co-expressed gene networks (modules) enriched for specific biological responses and transcription factor target genes that provide biological interpretation based on module induction or repression^19,20^. Induction or repression of each module, and by analog the associated biological processes annotated for that module, is quantified using a single eigengene score (EGs) based on the fold-change expression of individual gene network memberships^21^.

By combining our LCM-TempO-Seq approach with the Kidney TXG-MAPr platform, we identified different sensitivity in each nephron region towards cisplatin as well as distinct temporal response dynamics of gene network activities, with outer-medulla proximal tubules being the most affected region. While whole kidney-TempO-Seq lacked the ability to cover the spatial heterogeneity as well as sensitivity of the cellular responses. Moreover, we observed that the urinary injury biomarker clusterin most accurately reflects the sustained injury response in the outer-medulla region.

## Materials and Methods

### Animal experiments

#### Cisplatin dose-response experiment

Twenty male Sprague-Dawley rats (Crl: CD® [SD]; 8 to 9 weeks old; body weight 320-360 g) were pair-housed in type 3 cages with disposable inlays under controlled conditions (temperature: 20 to 24°C; humidity: 40 % to 70 %) and fed with standard food and water *ad libitum*. For urine collections, rats were kept individually in metabolic cages overnight under fasted conditions but with unrestricted access to water. To locate the fasted urine collection period in the resting period of the animals, the light cycle was inverted (dark phase: 7 AM - 7 PM, light phase: 7 PM – 7 AM) starting 4 days before dosing. The rats were randomly assigned to five groups (4 males/group) and received either the control vehicle (0.9% sodium chloride) or cisplatin (Teva, 1 mg/ml diluted with 0.9 % NaCl) at dosages of 1, 2.5, 5, or 7.5 mg/kg/dose via a single intravenous bolus administration in the tail vein. The blood (collected under isoflurane inhalation anesthesia from the sublingual vein) and urine were collected every day for biomarker identification. The rats were euthanatized by exsanguination under sevoflurane (Sevorane®, AbbVie Deutschland GmbH&Co. KG, Knollstraße, Ludwigshafen) anesthesia performed according to the German Federal Animal Welfare Act at collection days. Directly after the exsanguination, the right kidney was collected and fixed with 10 % formaldehyde for further embedding in a paraffin block for immunostaining and histopathological assessment. The other left kidney was snap frozen for tissue platinum determination and LCM method optimization.

#### Cisplatin time course experiment

Fifty-four male Sprague-Dawley rats (Crl: CD® [SD]; 8 to 9 weeks old; body weight 320-360 g) were randomly grouped into 18 groups (3 rat per group) with 4 control groups. The general housing conditions were the same as in the dose response experiment, including an inverted light cycle. As opposed to the first study, rats were single housed in Type 3 cages with disposable inlays for the first five days post dose to best protect personnel from cisplatin exposure. After these initial 5 days, the remaining rats were group housed in their dose groups (3 animals/cage) in 2000P type cages. Rats were administered with 5 mg/kg i.v. cisplatin as this dose was found to elicit clear modulation of cellular responses based on the dose-response study; control groups were received the vehicle alone (0.9% saline). The rats were sacrificed for designated time points (1, 2, 4 and 24 hours; 3, 5, 8, 10, 12, 15, 20 and 28 days). The control groups were sacrificed at 24 hours, 8, 15 and 28 days. Six rats grouped into 2 groups were treated as the reserve group (28 days cisplatin). Urine and blood from this study were collected from day 0-15, on day 20, and on day 28. For both studies, blood (200 µL) was collected from the sublingual vein under sevoflurane (Sevorane®, AbbVie Deutschland GmbH&Co. KG, Knollstraße, Ludwigshafen) anesthesia. Serum was immediately harvested from the blood and stored at -20°C for clinical chemistry purposes (BUN and creatinine). Urine was centrifuged to collect the supernatant for biomarker identification. Rats were euthanatized by exsanguination under sevoflurane (Sevorane®, AbbVie Deutschland GmbH&Co. KG, Knollstraße, Ludwigshafen) anesthesia performed according to the German Federal Animal Welfare Act at collection days. Directly after the exsanguination, the right kidney was collected and fixed with 10 % formaldehyde for further embedding in a paraffin block for immunostaining and histopathological assessment. The other left kidney was snap frozen for tissue platinum determination.

#### Biomarker measurement

The determination of kidney injury biomarkers was performed with commercial kits. Albumin and Lipocalin-2/NGAL values were obtained using MILLIPLEX MAP Rat Kidney Toxicity Magnetic Bead Panel 2 (Millipore Sigma, USA) with 1:500 dilution. KIM1 and clusterin values were obtained utilizing MILLIPLEX MAP Rat Kidney Toxicity Magnetic Bead Panel 1 (Millipore Sigma, USA) without dilution. Serum biochemical profiles (BUN and creatinine) were evaluated with a cobas c501 (Roche Diagnostics GmbH, Mannheim, Deutschland). All the steps were performed according to the standard protocol from the manufacturers.

#### Laser Capture Microdissected samples collection

Paraffin blocks were sectioned with 8 µm thickness and mounted on PEN membrane slides (Zeiss, 415190-9081) and thereafter stained with fast haematoxylin-eosin staining. The slides were deparaffinized with xylene twice and rehydrated by dipping the slides in 100% EtOH twice, 70% EtOH, 50% EtOH, and washed with RNAse free water. Slides were further stained with hematoxyline gill (Sigma, GHS232) and washed with RNAse free water followed by washing the slides with RNAse free PBS for 30 seconds. Slides were then washed with RNAse free water and stained with eosin (Merck, 102439) twice and dehydrated with 100% EtOH 2 times each 30 seconds. The duration of every step unless differently mentioned was 1 minute. Stained slides were directly subjected to the LCM steps.

The LCM was performed using P.A.L.M MicroLaser Technology with a pulsed UV laser (wavelength: 337 nm) connected to a Zeiss Axiovert 200 microscope and controlled using PALM® RoboSoftware (version 4.6). Three different nephron segments were dissected: glomeruli (GM), cortical proximal tubules (CPT) – the proximal tubules located in the cortex of kidneys, and outer- medulla proximal tubules (OMPT) – the proximal tubules located in the outer stripes of the renal medulla. In total 200.000 – 400.000 µm^2^ of surface area LCM samples were collected in the adhesive cap of a collection tube (Zeiss, 415190-9201) followed by storage at -80°C until sequencing. For each sample, we repeated the collection process twice (technical replicates). For each treatment, single whole kidney FFPE sections were also collected for sequencing. The procedures of the RNA collection and sequencing process are explained elsewhere^10^.

For the frozen sections, the kidneys were snap-frozen in liquid nitrogen pre-cooled isopentane. The frozen kidneys were embedded with Tissue-TEK O.C.T (Sakura) at -80°C in a cryomold. The kidneys were then sectioned with 30 µm thickness and mounted in the super frost slides (Fisher scientific).

#### Immunohistochemistry

Four µm kidney sections mounted on super frost slides (Fisher scientific) were dried overnight at 37°C. After deparaffinization and rehydration process (twice xylene followed by 100% EtOH twice, 90% EtOH, 70% EtOH, and twice water for 5 min each) slides were incubated with 3% of H2O2 in PBST_0.05_ for 20 minutes to deplete the activity of endogenous peroxidases. Antigen retrieval steps were performed by boiling the slides in 0.01 M citrate buffer pH 6.0 in PBST0_.05_ for 15 minutes at 120 °C with an autoclave. After 20 min cooling sections and blocking with 3 % w/v BSA-PBST0_.05_ for 1 hour at room temperature, slides were incubated overnight at 4°C with the desired diluted antibody solution (Table 1) in 0.5% BSA- PBST0_.05_. Then slides were washed with PBST0_.05_ for 10 min and the 1 hour secondary antibody incubation was performed at room temperature (1:100 dilution of antibody of choice in 0.5 % w/v BSA- PBST0_.05_). Slides were then washed with PBST0_.05_ for 10 min followed by addition of DAB solution (Thermo Fisher, 34002) for 10 minutes and subsequent twice 3 min wash with PBST0_.05_. Then slides were counterstained with haematoxylin (Sigma Aldrich – GHS232) and dehydrated with reverse EtOH sequences and xylene. A glass coverslip was mounted to the slides with Entellan (Merck, 107960) and the slides were stored at room temperature. Immunostained slides were further scanned with 3DHISTECHT digital slide scanner and all the images were processed with SlideViewer 2.5 and ImageJ 1.53c. The positive nuclear count for p53-pSes15 and Ki67 were quantified manually by counting the percentage of DAB-positive nuclei. For hematoxylin/eosin (H&E) staining slides were deparaffinized as above followed by hematoxylin (Sigma, GHS232) solution for 10 minutes followed tap water for 15 minutes and eosin (Merck, 102439) staining for 10 minutes followed by the dehydration and mounting as above.

**Table 1.** Overview of the *in vivo* experiments

| *Experimental series* | *Intervention* | *Sample size* | *Outcomes measures* |
| --- | --- | --- | --- |
| 1 | Cisplatin dose 0, 1, 2.5, 5, 7.5 mg/kg (iv SD) with follow-up 72 hours post dose | 20 animals, 4 animals per group | Transcriptomic profiles in the kidneys, immunohistochemistry in the kidneys, platinum content in the kidneys, urine biomarkers (Multiplex NGAL, KIM-1, albumin, osteopontin, Rat clusterin (singleplex), Alpha GST and glucose), and serum biomarkers (creatinine and BUN) |
| 2 | Cisplatin dose 5 mg/kg (iv SD) with follow-up 1 hours, 2 hours, 4 hours, 1 day, 3 days, 5 days, 8 days, 10 days, 12 days, 15 days, 20 days, 28 days post dose. Control animals were taken on 1 day, 8 days, 15 days, and 29 days. | 48 animals, 3 animals per group | Transcriptomic profiles in the kidneys, histopathology in the kidneys, immunohistochemistry in the kidneys, platinum content in the kidneys, plasma (toxicokinetics), urine biomarkers (KIM1 alpha-GST, clusterin, albumin, glucose, NGAL (lipocalin-2), urine creatinine and osteopontin (OPN)), and serum biomarkers (*in vivo*: creatinine and BUN, at necropsy full clinical chemistry) |

**Table 2.** List of primary and secondary antibody used for the immunostaining experiments.

| Protein | Host | Ab catalogue | Dilution | Final concentration |
| --- | --- | --- | --- | --- |
| KIM1 | Goat | R&D Systems (AF3689) | 1:15 | 15 µg/ml |
| Clusterin | Mouse | R&D Systems (AF2747) | 1:15 | 13 µg/ml |
| TP53-S15 | Rabbit | CST (9284) | 1:15 | Initial concentration not provided |
| Vimentin | Mouse | Abcam (ab8069) | 1:300 | 3.3 µg/ml |
| Ki67 | Rabbit | Abcam (ab16667) | 1:150 | 4.8 µg/ml |
| Anti-goat | Donkey | Jackson (705-035-003) | 1:100 | 8 µg/ml |
| Anti-mouse | Goat | Jackson (115-035-003) | 1:100 | 8 µg/ml |
| Anti-rabbit | Goat | Jackson (111-035-003) | 1:100 | 8 µg/ml |

#### Inductively Coupled Plasma-Mass-Spectrometry (ICP-MS) for platinum (Pt195) measurement

Frozen kidneys were embedded in tissue-tek O.C.T. (Sakura) and solidified at -30°C in the cryo-chamber. Kidneys were sectioned with 30 µm thickness and mounted on super frost slides (Fisher scientific). Sections were dried at room temperature for one hour and scraped to an eppendorf tube (4 sections per condition ∼100 µm thickness) containing lysis buffer (20 mM Tris-HCl, pH 7.4, 137 mM NaCl, 2 mM EDTA, 1% Triton, 10% glycerol). The lysate was 10x diluted in 65% HNO3 and heated at 95°C for 1 hour. For plasma samples, the plasma was diluted 10x with 65% HNO3 (SuprapurR, Merck) and heated at 95°C for 1 hour. The nitric acid mixture from both kidney lysate and plasma were cooled down at room temperature and diluted 10x with demineralized water. This solution was then subjected to the Pt measurement with the ICP-MS instrumentation. (NIST)-traceable 1000 mg/L elemental standards were used (TraceCERTR, Fluka) for preparation of calibration and internal standards. Five trace elemental calibration standards for ICP-MS analysis were prepared using National Institute of Standards and Technology NISTtraceable 1000 mg/L Pt standard: 0, 1, 5, 20, 100 ug/L. 10 ug/L Rh and in were used as internal standard. The standards and samples were analyzed for trace elements using the NexIONR 2000 (PerkinElmer) ICP-MS equipped with a concentric glass nebulizer and peltier-cooled glass spray chamber. An SC2 DX autosampler (PerkinElmer) was connected to the ICP-MS for sample introduction. Syngistix™ Software for ICP-MS (v.2.5, PerkinElmer) was used for all data recording and processing. The final platinum content was calculated from the ICP-MS output with the correction from the dilution factor. The tissue Pt content was normalized with the total protein concentration determined with the BCA assay (Thermo Fisher, 23225).

#### TempO-Seq gene expression analysis of cisplatin-exposed kidneys

Firstly, we evaluated the most optimal preservation method of the kidneys (FFPE and snap-frozen) for the TempO-seq using the whole rat transcriptome consisting of >20000 probes. We compared the %mapped reads (the percentage of the correctly mapped reads of the genes during the sequencing processes) of the kidney sections preserved with these two approaches. The preservation method resulting higher %mapped read was chosen for the further LCM-sample collection steps. Upon retrieving the sequencing outcomes, probe-alignment steps were performed according to the pipeline from BioSpyder^22^. The read count values of correctly aligned genes were retrieved as the raw count values. Samples with lower than 10 % of %mapped reads were excluded (Suppl. figure 1B and 2A, dose-response and time response experiment). Raw read count values were aggregated from the technical replicates. The summed raw read count value was transformed by adding 1 to every count. The probes which had base mean lower than the first quantile of base mean value were excluded. Correlation analysis was performed subsequently to check the similarity of the two replicates in each kidney samples; samples with lower than 0.8 Pearson correlation values were excluded from further analysis.

Count per million (CPM) normalization was performed on the quality-controlled data and the value of log2 fold change to the control (the non-exposed samples from the same segments) of the normalized count was calculated using DESeq package^23^. The differential expressed genes (DEGs) were defined by the genes passing the threshold of adjuster p-value < 0.05 and log2 fold change values > |2|. The replicate correlation and normalized count distribution plots are depicted in Suppl. figure 1C and 2B for dose-response and time-response experiment respectively. For the temporal response experiment the controls from different time points were combined as the control samples from all the time points showed no transcriptional differences (Suppl. figure 3A). To qualify and quantify modulation of the gene network (modules) representing the cellular responses, gene log2 fold change values were uploaded to the Kidney TXG-MAPr platform (manuscript in preparation). The log2 fold change values are tabulated in Suppl. table 1 and 2 for dose-response and time dynamics experiments, respectively. Module scores were expressed based on module eigengene score (EGs) calculation derived from the log2FC expression value of every gene membership^20^ (tabulated in Suppl. table 3 and 4 for dose-response and time dynamics experiment, respectively). The significant modules were selected based on these criteria : 1. The modules have >7 gene memberships; 2. The coverage of the modules from the external datasets is >50%; 3. The maximum EGs of the modules is >2. The module memberships of all the important modules (modules that are mentioned in the study – linked to the mechanisms of cisplatin induced kidney injury with clear dose-response and time-response patterns) are listed in Suppl. table 5. The annotation of the selected modules are listed in Suppl. table 6. To evaluate the expression of marker genes in the specific nephron segments, we used the list of the marker genes obtained from published single cell RNA sequencing analysis of mouse kidney^24^ (Suppl. table 7). We used the rat orthologue list from the rat genome database (RGD)^25^ to match the orthologues of rat genes to mouse genes. The log CPM normalized values were used to evaluate the expression of the marker genes in each segments.

#### Data Representation

Correlation analysis was performed with the R function from the package Hmisc^26^. Statistical analysis (independent student t-test) was performed using GraphPad Prism 8. The hierarchical clustering was performed with “*Ward D2*” algorithm applied to the Euclidian distance between measured variables. Illustrator 22.1 and R^27^ (ggplot2^28^ and pheatmap^29^) were used for graphical representation of the results.

## Results

### Targeted RNA sequencing of micro-dissected nephron regions reveals coherent segment specific signatures

The aim of our study was to investigate mechanisms of cisplatin-induced tubular injury based on spatio-temporal transcriptional responses in different nephron segments. First, we established an LCM methodology and transcriptional analysis strategy to assess the transcriptomes in different nephron segments using targeted RNAseq, TempO-Seq. We collected glomeruli (GM), cortical proximal tubules (CPT) and outer-medulla proximal tubules (OMPT). The FFPE samples exhibited higher RNA quality compared to frozen samples resulting in higher %mapped reads (Suppl. figure 1A). With this preservation method, we proceeded to collect the nephron segments using LCM methodology. The RNAseq results indicated that the clustered of samples with %mapped reads >10% (Suppl. figure 1B), showed high replicate correlation, and similar distribution of normalized counts, except for GM (Suppl. figure 1C). A Principal Component Analysis (PCA) indicated a unique gene expression pattern for whole kidney, GM, CPT and OMPT (Figure 1B). Moreover, characterization of the segment specific gene signatures obtained from the single cell atlas of mouse kidney^24^ (Suppl. table 7) validated the LCM-mediated nephron segment collection (Figure 1C). CPT samples showed the expression pattern of the tubular markers from the segment 1 and segment 2 of the proximal tubules. OMPT samples displayed an expression profile of the marker genes of the proximal tubule from segment 2 and segment 3. Podocyte markers were enriched in GM samples. We further evaluated the expression of well-established regional nephron marker genes in the transcriptomics data (*Nphs1*/podocytes; *Slc5a2*/proximal tubule s1 and s2; *Slc22a2*/proximal tubule s3; *Kcjn1*/distal tubule; *cldn8*/loop of Henley; *Slc8a1*/collecting duct)^30,31^. The expression of *Nphs1* was found the highest in GM samples. *Slc5a2* and *Slc22a2* were enriched in the CPT and OMPT samples, respectively (Figure 1D). While all marker genes were expressed in the whole kidney samples, the distal tubule, loop of Henle and collecting duct markers, *Kcjn1, cldn8* and *Slc8a1, respectively*, were far less abundant in CPT, OMPT and GM. Altogether, the data provided confidence that our LCM-based, spatially-anchored transcriptomics analysis allows for a segment-specific analysis of transcriptional responses.

**Figure 1.**
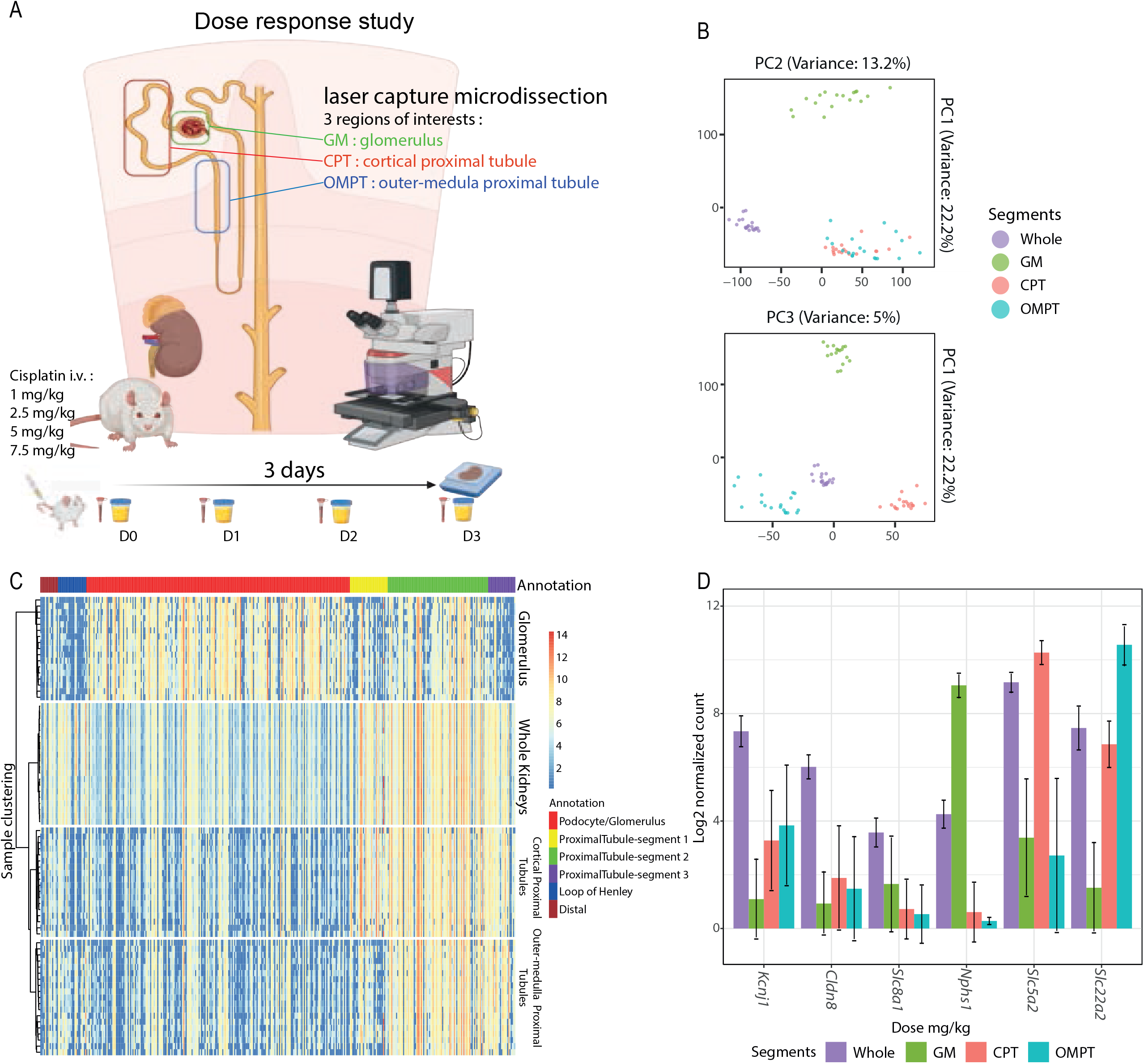
Transcriptomics analysis of nephron segments reveals coherent cell type specific signature. Overview of the 3 days dose response *in vivo* experiment during which blood and urines were samples. After sacrifice, tissue samples ofspecific tissue regions were collected with laser capture microdissection (LCM) technology from FFPE embedded kidneys. This figure was created with BioRender.com. (B) PCA plots of each individual collected samples made from gene expression values (Whole kidney, GM, CPT, and OMPT) displayed both PC2 and PC3. (C) Heatmap showing the gene expression profiles of all tissue samples of specific marker genes for particular nephron segments^24^. The sample clustering was performed using ‘Ward D2’ algorithm. The color on the annotation bar indicates the group of the marker genes expressed in the specific nephron segments that was extracted from the previous study^24^. The color scale of the heatmap indicates the log2 normalized count of each gene. (D) Gene expression value of specific nephron region markers (*Kcjn1* : distal tubule, *cldn8* : loop of henley, *Slc8a1* : collecting duct, *Nphs1* : podocyte/glomerulus, *Slc5a2* : proximal tubule s1 and s2, *Slc22a2* : proximal tubule s3). Sample types are depicted with the colors of the plots.

### Outer-medulla proximal tubules are the most susceptible region to cisplatin-induced nephrotoxicity

Having established the LCM approach, we next determined the most optimal dosing regimen for the temporal segment specific transcriptional responses in the kidney. For this purpose we performed a dose-response study to assess the segment-specific transcriptional responses 72 h after cisplatin treatment. The number of differentially expressed genes (DEGs) in whole kidneys and the nephron segments displayed a clear dose-response relationship to cisplatin exposure with OMPT showing the highest number at the highest dose followed by the glomerulus samples (Figure 2A). We then leveraged kidney TXG-MAPr to gain insight into the cisplatin responses based on changes in module-based responses. We found that 90% of the modules showed more than 75% coverage with our kidney TempO-Seq data (Suppl. figure 3B). The high module coverage ensured high representation towards the module EGs, thereby increasing the accuracy of the biological interpretation of transcriptomics data. Consistent with the number of DEGs, the number of significantly (de)activated modules exhibited a dose-response trend in the OMPT and whole kidney, which was less pronounced in GM and CPT (Figure 2B and 2C, Suppl. Table 3). Interestingly, number of significant modules in whole kidneys showed the lowest number indicating that at the whole kidney perspective, the activation of cellular responses was attenuated. Despite the high number of DEGs and significant modules in GM, the annotation of the modulated gene networks shows non-specific and non-dose dependent cellular responses (Figure 2C-inset 1a and b, and Suppl. table 8). While the toxicogenomic map of the OMPT exhibited stronger and broader module responses compared to whole kidneys (Figure 2C-inset 2, 2E).

**Figure 2.**
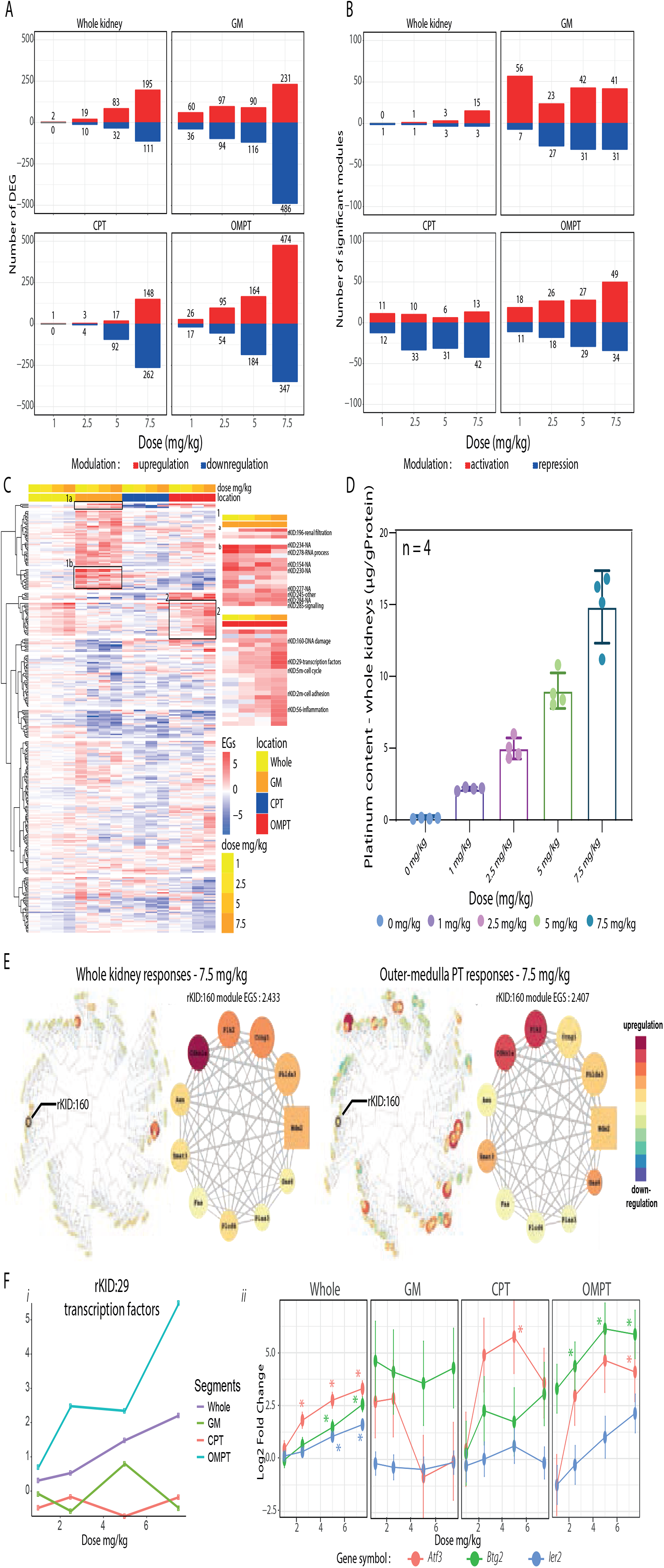
Dose response effects of spatially-anchored transcriptional changes induced by cisplatin. (A) Numbers of Differentially expressed genes of each sample types (whole kidney, glomerulus, CPT, and OMPT – threshold : adj p-val < 0.05, log2 fold change > [2]). Red bars indicate upregulation of the genes and blue bars indicate downregulation of the genes. (B) Numbers of significantly de(activated) modules of each sample types (whole kidney, glomerulus, CPT, and OMPT – thresholds : gene memberships >7, module coverage >50%, and EGs > |2|). Red bars indicate activated modules and blue bars indicate repressed modules. (C) Heatmap showing an overview of the kidney TGX-Mapr module (n=4) activities. The dose bar indicates the cisplatin dose and the segment bar indicates the sample type. Each block in the heatmap represents a response of a module in a particular sample type at the specific dose. The color of the heatmap indicates the magnitude of the module responses (eigengene scores; red: activation, blue: deactivation). Inset 1a and b contain the modules highly modulated in GM and inset 2 contains the modules highly modulated in OMPT. (D) Platinum (Pt 195) content measured in kidney tissue with ICP-MS. The color of the boxplot represents the cisplatin dose with each dot representing the value of each sample in the same condition (biological replicates). (E) Toxicogenomics maps of whole kidneys and OMPTs for dose cisplatin 7.5 mg/kg (left). Each circle on the map represents one module (highlighted: rKID:160-DNA damage module), color and the size of the circle indicate the magnitude of the response modulation (red: activation, blue: deactivation). Detailed overview of module WGCNA |Kidney:160 membership (right). Color and the size of the nodes indicate the magnitude of the response modulation (red: activation, blue: deactivation), the square node is the hub gene of the module (ii). (F) Dose response trend of rKID:29-transcription factors module activity (eigengene score) in each nephron region and whole kidneys. The color of the plot represents the sample type (i). The dose response plots of 3 highest correlated gene members of rKID:29 in each sample type. The color of the plots show the expression of gene memberships (error bars represent SEM). The stars represent the significant log2 fold change values (adj p-val < 0.05) (ii).

The dose-response pattern of the toxicogenomics responses of the kidneys matched the platinum content (Figure 2D). The cisplatin kidney burden was associated with increasing transcriptomics perturbation and more severe (proximal) tubular injury (Suppl. figure 4B), particularly in the OMPT region possibly due to higher Pt accumulation. Overall, these results indicate that transcriptional changes in injured OMPT was enhanced that higher cisplatin doses in male rats.

### Spatially-anchored transcriptional changes reveal mechanistic details for cisplatin injury and repair

Next, using kidney TXG-MAPr we explored and quantified the activation of segment-specific biological perturbations that are associated with cisplatin-induced nephrotoxicity. One of the pronounced responses was activation of module rKID:29 containing transcription factors. Only OMPT clearly exhibited strong activation of this module suggesting that OMPT is the only region contributing its activation captured in whole kidneys (Figure 2Ei). Interestingly, the top 3 highest correlated gene memberships, *Btg2, Atf3*, and *Ier2* are known to be the transcription factors linked to the early response of cellular injury leading to various activation of cellular responses^32–34^ (Figure 2Eii). Only whole kidney, CPT, and OMPT exhibited clear dose-response upregulation of *Btg2* and *Atf3*, while the upregulation of *Ier2* was only found in OMPT and whole kidney. Lastly, *Btg2* and *Atf3* were non-significantly upregulated in GM.

Furthermore, we focused on four cellular responses: i) DNA damage response, ii) mitochondrial and energy production, iii) actin cytoskeleton and iv) immune response (Figure 3A and 3B). Activation of the DNA damage response is the primary cellular stress response towards cisplatin^35,36^. Among all modules, rKID:160 (Figure 2C) had the highest enrichment for GO:BP terms related to p53 and the highest enrichment for p53 target genes including *Cdkn1a* and *Plk2*^37,38^. While no difference was observed in activation of rKID:160 between whole kidney, CPT and OMPT at 7.5 mg/kg, this module was already activated at the lowest cisplatin dose in OMPT (Figure 3Ai). Five strongest upregulated genes involved in cell cycle arrest resembled the dose-response pattern of rKID:160 (Figure 3Aii).

**Figure 3.**
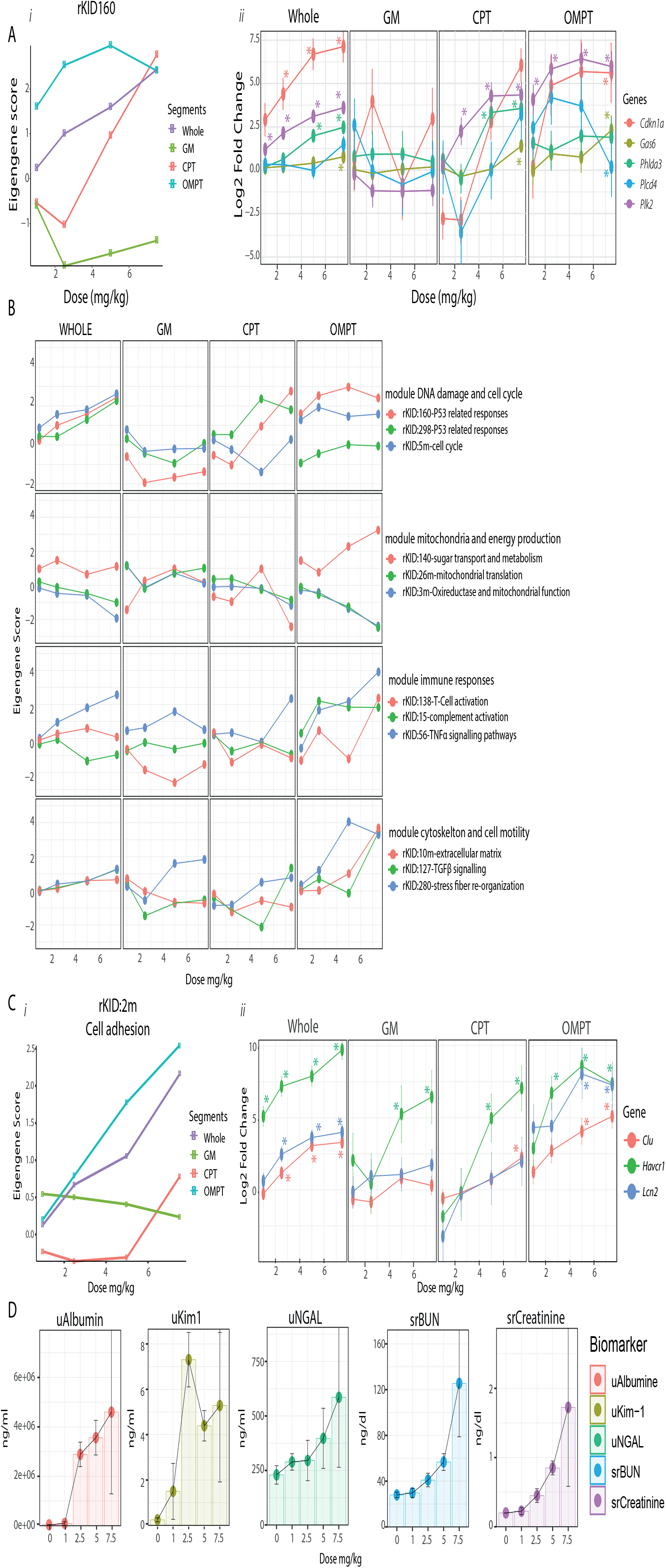
Dose response activation of cellular responses elicited in different nephron segments. (A) Dose response trend of rKID:160-DNA damage module activity (eigengene score) in each nephron region and whole kidneys. The color of the plot represents the sample type (i). The dose response plots of 5 highest expressed gene members of rKID:160 in each sample type. The color of the plots show the expression of gene memberships (error bars represent SEM). The stars represent the significant log2 fold change values (adj p-val < 0.05) (ii). (B) Cellular responses elicited in the nephron segments. The responses are grouped into 4 different injury responses of cisplatin: DNA damage (i), mitochondrial function (ii), cytoskeletal (iii), and immune responses (iv). Each response is represented by 3 different modules. The color of the plot represents the activity of each module. (B) The dose-response plot of rKID:2m-Biomarker module upon cisplatin exposure. The color of the plots represents the response of each segment (Bi). Dose-response plots of the gene expression level of 3 kidney biomarkers from rKID:2m (Clusterin, KIM1, Lcn2). The error bars display the SEM. The stars represent the significant log2 fold change values (adj p-val < 0.05) (Bii). (C) Serum and urinary biomarkers upon cisplatin exposure measured after 72 hours (u - urine, sr - serum). Error bars indicate SD.

Yet, rKID:298 (another P53 related-response) was particularly activated in CPT, associated with stronger upregulation of *Mgmt* and *Nudt11 (*rKID:298 membership) (Suppl. figure 5A) known to protect cells from chemical stress^39–42^. The activity of cell cycle responses represented by rKID:5m displayed a high activity in OMPT and whole kidney(Figure 3B). With respect to the energy metabolism represented by modules rKID:140 (sugar transport and metabolism), rKID:26m (mitochondrial translation), and rKID:3m (oxireductase and mitochondrial function), OMPT seemed to be the most responsive. While the activities of rKID:26m and rKID:3m were repressed, the activity of rKID:140 increased at high doses, possibly leading to metabolic switch from mitochondrial metabolism to enhanced glycolysis as a result of direct mitochondrial damage by cisplatin^43^ and alternatively due to initiation of a regeneration program. Further, we interrogated cisplatin-induced inflammatory responses^44–46^ by selecting three inflammation-related modules: rKID:138 (T-cell activation), rKID:15 (complement activation), and rKID:56 (TNFα signalling). Despite moderate activation of the TNFα signalling pathway in CPT, it is much stronger in OMPT. On the other hand, the complement-mediated inflammation and T-cell activation were only induced in OMPT, potentially resulted from the strong activation of the TNFα signalling (rKID:56)^47,48^. Finally, we charted the activity of three cytoskeletal and cell motility-related modules: rKID:10m (extracellular matrix), rKID:127 (Cell migration-TGFβ signalling), and rKID:280 (stress fibre re-organization). Cisplatin-induced actin cytoskeleton reorganization was associated with tissue remodelling during severe kidney injury^49,50^. With increasing cisplatin doses, actin stress fibres are formed significantly in OMPT while the extracellular matrix is actively remodelled^49^. In addition, TGFβ signalling which is a key cue for both actin and ECM re- organisation leading to the increase of cell motility^51^ was initiated at 5 mg/kg also in OMPT. In summary, our data reveal the activation of various DIKI-related mechanisms in both CPT and OMPT. While the activation of cellular responses linked to the mechanisms of cisplatin induced kidney injury in GM was minimal.

Another cytoskeletal-related module, rKID:2m (cell adhesion) interestingly contains various genes typically described as relevant renal biomarkers, including *Havcr1*/KIM1, *Clu*/Clusterin, and *Lcn2*/Lipocalin2/NGAL^18^. This might suggest that the cells undergoing cytoskeletal re-organization also express the injury biomarkers. Cisplatin caused a dose-dependent activation of rKID:2m in whole kidney, CPT, and OMPT, with activation in OMPT and whole kidney at 2.5 mg/kg (Figure 3Ci). The upregulation of *Hacvr1* was found in each segment, confirmed by KIM1 immunostaining of kidney sections (Suppl. figure 5B). In contrast, the enhanced expression of *Lcn2* and *Clu* was more specific to the OMPT (Figure 3Cii). At day 3, the urinary albumin, KIM1 and NGAL showed a dose-dependent elevation, with KIM1 being the most sensitive and already elevated at lower doses and earlier time points, yet preceding overall loss of renal function based on serum creatinine and BUN analysis (Figure 3D and Suppl. figure 5C).

### Sustained spatial DNA damage drives enhanced renal injury responses specifically in OMPT

Next, we systematically mapped the spatio-temporal dynamics of cisplatin-induced renal injury responses. Based on our dose response study, we selected 5 mg/kg cisplatin and determined the transcriptional changes in whole kidney, CPT and OMPT at different time points up to 28 days (Figure 4A). GM was excluded given the non-dose dependent responses, absent of DNA damage, and low count for most of the genes based on the dose-response study. Overall, higher numbers of DEGs and significant modules were observed in OMPT compared to whole kidney and CPT, and increased drastically in the first three days, and persisted up to day 20 (Figure 4Bi and ii). The transcriptional time course profile was in line with the histopathological analysis, showing early necrosis peaking at day 5, followed by regeneration/degeneration and fibrosis which persisted throughout the time course (Suppl. figure 6A). Various cellular responses were modulated in OMPT in concordance with whole kidney responses with much higher module EGs (Figure 4C). Sustained increase in activity was observed for the modules that reflect DNA damage response, cytoskeletal remodelling, inflammation, and metabolism; the modules reflecting mitochondrial function were persistently repressed over time. Interestingly, in contrast to the sustained module activity in OMPT, the platinum levels rapidly reached the Cmax at 1 hour in the kidneys. Plasma platinum levels readily declined within 4 hours (T1_/2_ plasma <1 hour), while renal platinum levels remained relatively stable over the time course of the experiment (T1_/2_ kidney >10 days) indicative of persistent tissue binding (Figure 4D).

**Figure 4.**
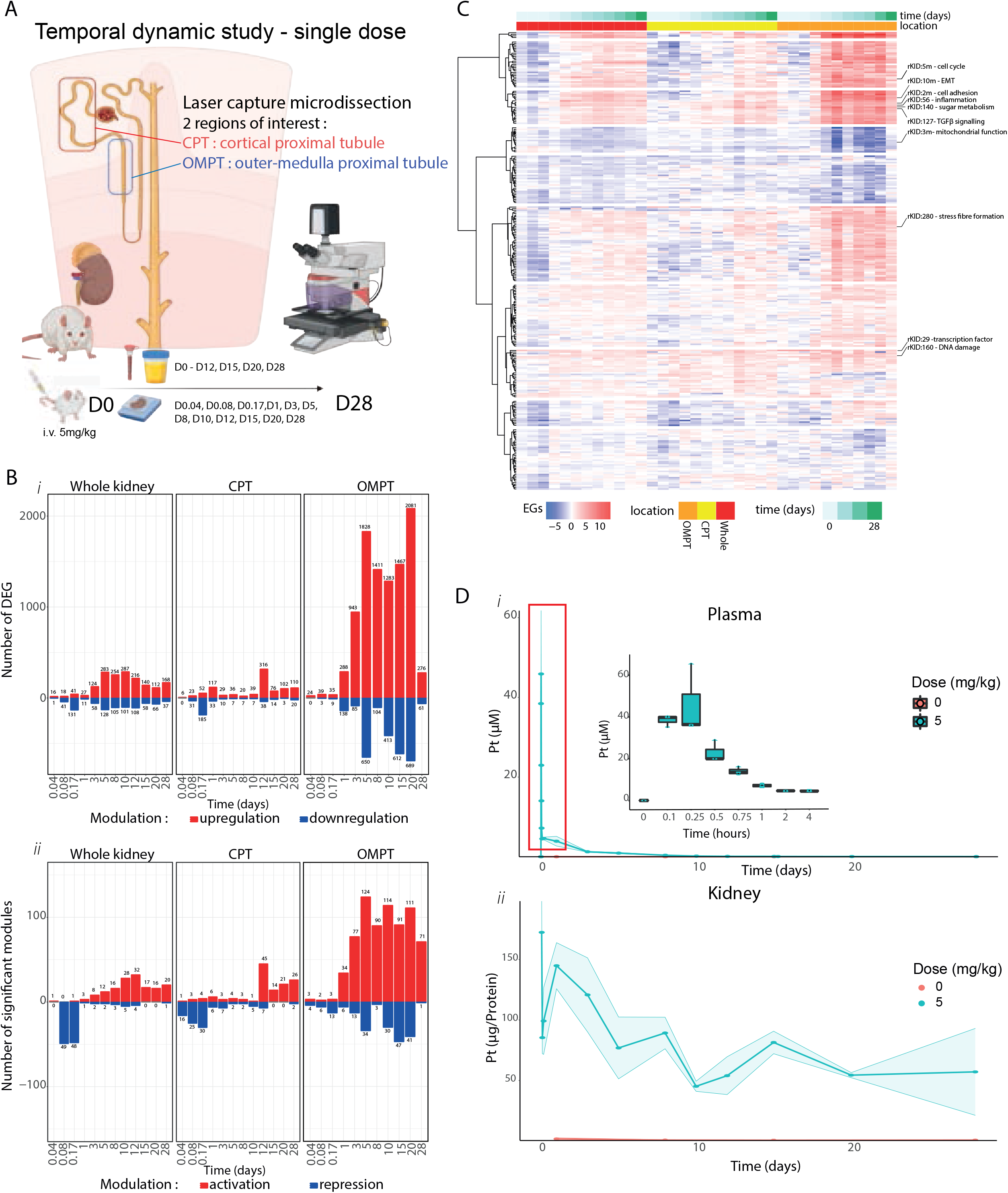
Temporal cisplatin transcriptional responses in whole kidney, CPT and OMPT. (A) Overview of the time dynamic experiment and sample collection timeline. Red tubes indicate blood collection, yellow jars indicate urine collection, and FFPE block indicate kidney collection. This figure was created with BioRender.com. (B) The number of differentially expressed genes (DEGs) of whole kidneys, CPT, and OMPT at each specific time points – threshold : adj p-val < 0.05, log2 fold change > [2]). Red bars indicate the number of upregulated genes and blue bars indicate downregulated genes (i). Numbers of significantly de(activated)modules of each sample types (whole kidney, CPT, and OMPT – thresholds : gene memberships >7, module coverage >50%, and EGs > |2|). Red bars indicate activated modules and blue bars indicate repressed modules (ii). (C) Heatmap showing the overview of the temporal dynamics of the module activity. The time legend indicates the duration of cisplatin exposure, and the segment legend indicates the nephron regions. Each block in the heatmap represents a response to a module in a particular sample type at a specific time point. The color of the heatmap indicates the magnitude of the module responses (eigengene scores; red : activation, blue : deactivation). (D) The plots of platinum (Pt 195) content measured in plasma (top, panel i) and kidney tissue (bottom, panel ii). The color of the plot represents the administered dose : 0 mg/kg (red) and 5 mg/kg (blue) and the shadow of the plot represents the standard deviation values. The inset in the plasma Pt content plot indicates the detailed overview of the Pt content at the early time points (0-4 hours). The error bar represents the SD.

Given the sustained platinum levels in the kidney, we wondered which nephron segment would be impacted most by cisplatin. On day 5, the kidney TXG-MAPr clearly demonstrated strong overall module activation in OMPT (Figure 5A) reflected the same modules as in the third day from the dose response study yet with higher activity (Figure 2C). Despite that CPT showed lower module activities on day 5. The DNA damage response (rKID:160), the initiating event in relation to cisplatin injury, was equally strongly activated on day 1 in both OMPT and CPT. Thereafter its activity declined in CPT, but remained high till day 12 in OMPT (Figure 5Bi). Similar to the parent module, the top 5 highest upregulated gene memberships showed the strongest responses in OMPT where they remained elevated up to 28 days, except for *Plk2* which had comparable induction in CPT and OMPT (Figure 5Bii).

**Figure 5.**
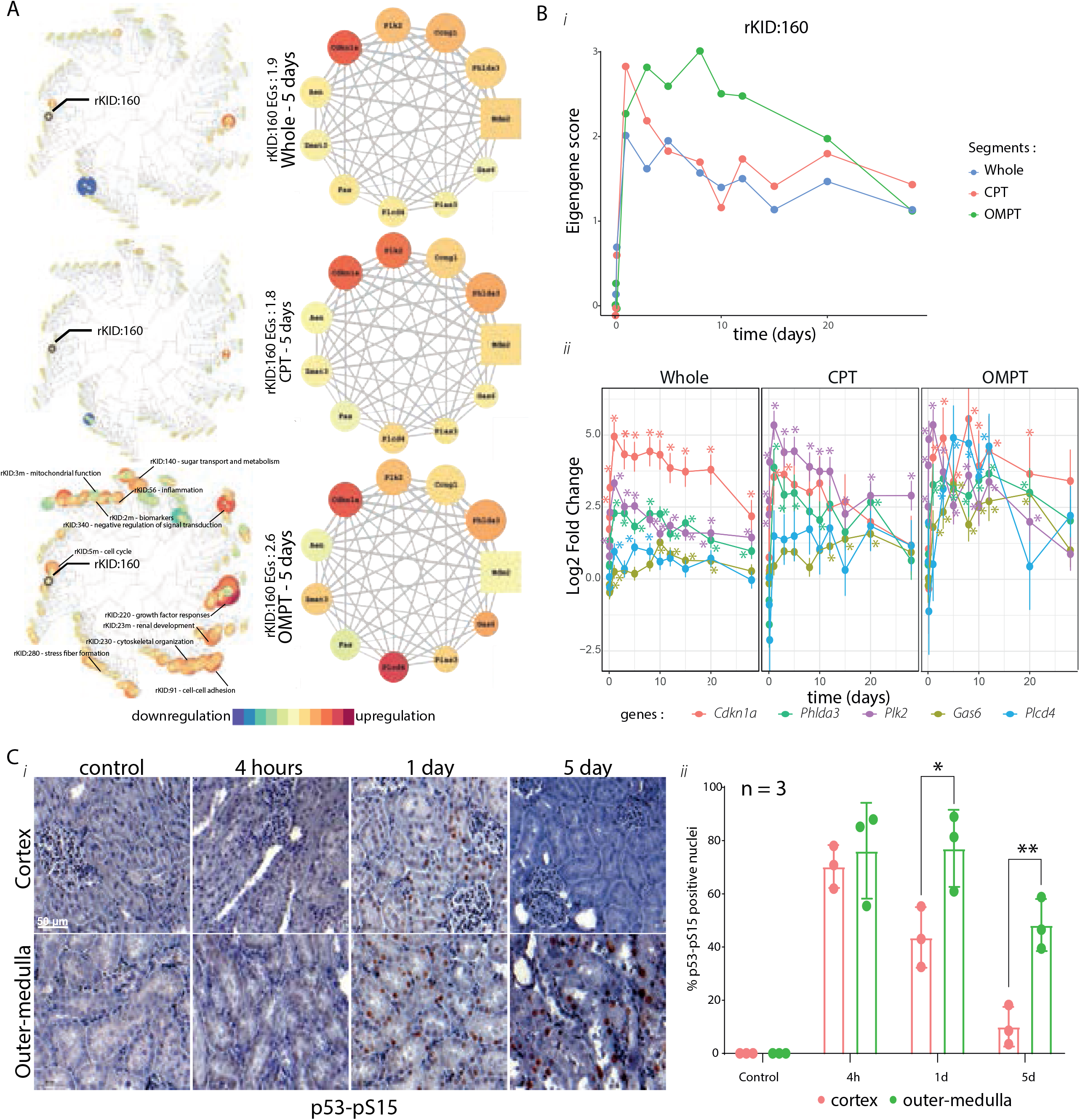
Assessment of the temporal cisplatin-induced DNA damage response in CPT and OMPT. (A) Toxicogenomics maps of cellular responses of the whole kidney, CPT, and OMPT assessed at day 5 of exposure. Each node on the map represents one module (highlighted: rKID:160-DNA damage). (B) Modulation of rKID:160 memberships in the whole kidney, CPT, and OMPT samples on day 5 after cisplatin exposure. The color and the size of the node indicate the activity of the responses (red: activation, blue: deactivation), the square node is the hub gene of the module. (C) Temporal dynamic plots of rKID:160 – DNA damage in the whole kidney (blue), CPT (red), and OMPT (green) (i). The time dynamic plots of 5 highest upregulated (in OMPT) module memberships of rKID:160-DNA damage in each sample type (ii). The color of the plots indicates the expression of gene memberships (error bars represent SEM). The stars represent the significant log2 fold change values (adj p-val < 0.05). (D) Nuclear staining of phosphorylated P53 protein - serine 15 in the cortex (left) and outer-medulla (right) – 60x magnification (i). The quantification of p53-pS15 positive nuclei in the cortex (red) and outer-medulla area (green) (ii). The error bar indicates the standard deviation values. Each dot in the plot shows the value of replicate. The stars indicate the p-value derived from independent student t-test indicating the significant comparison between cortex and outer-medulla regions (* p-val <0.05 – 0,01, ** p-val <0.01 – 0.001, p-val *** < 0.001) (ii).

To support our transcriptomics outcomes, we also performed the immunostaining of phosphorylated p53-Ser15 (p53-pS15) protein as a marker of the DNA damage response^52^ (Figure 5Ci). The phosphorylation of p53-Ser15 increased within 4 hours of cisplatin administration both in the tubules of the cortical and outer medulla regions with around 70% positive nuclei (Figure 5Cii). This was in agreement with the transcriptomics data, which revealed a rapid onset of the DNA damage response in both CPT and OMPT but a decline in the CPT. The p53-pS15 staining of tubular nuclei was sustained and increased in intensity only in the outer-medulla region, but declined in cortical region (Figure 5B and C).

Next, we explored the temporal relationship between the spatial anchoring of the DNA damage response and the modules that reflect critical DIKI-related cellular responses identified in the dose response study (Figure 6 relative to Figure 3A). Firstly, the TNFα signalling pathway represented by rKID:56 – *Icam1* (NF-ĸB downstream response), was rapidly elicited in OMPT almost in parallel to the DNA damage response, with limited activity in CPT; the inflammation response peaked between day 3 and 10 and then remained relatively high till day 28. The TGFβ signalling (rKID:280 – *Tgfbr2* (TGFβ receptor 2)) also exhibited a rapid activation in OMPT and later in whole kidneys and CPT. Early onset of the disruption of oxireductase and mitochondrial functions and induction of sugar transport and metabolism, represented by rKID:3m – *Mpc2* (mitochondrial pyruvate carrier 2) and rKID:140 – *Pfkp* (glycolysis regulation), respectively, was observed in OMPT. Interestingly, the temporal activation of cytosolic glucose metabolism was inversely correlated with a decline of mitochondrial function and paralleled the DNA damage response. These dynamics were not observed in CPT. Altogether, the sustained activation of the DNA damage responses in particularly OMPT is linked to the rapid activation of DIKI-related cellular injury responses. These spatial response dynamics relationships could not be identified from the whole kidney transcriptome data.

**Figure 6.**
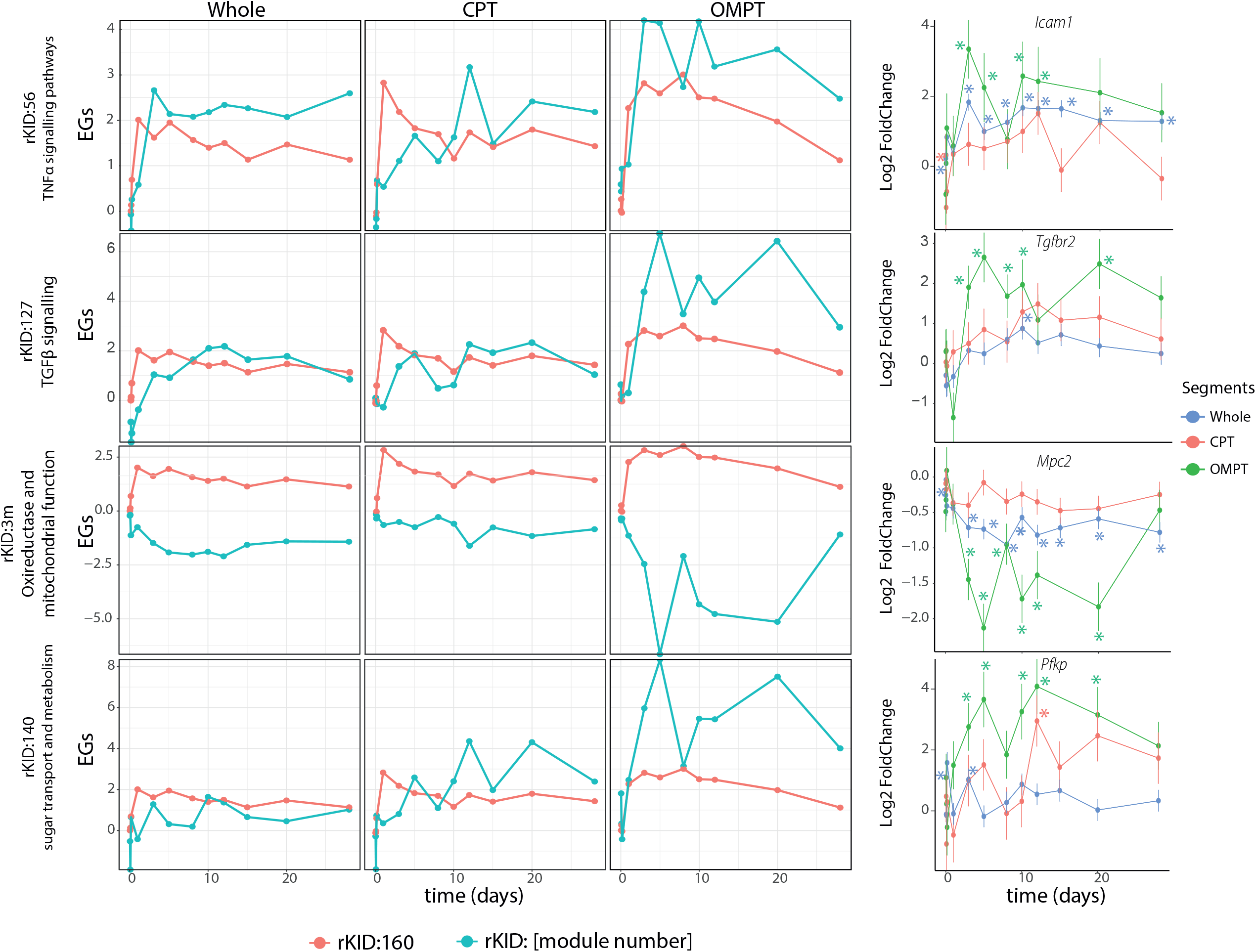
Comparative analysis of the temporal progressive cisplatin renal injury response in CPT and OMPT. Temporal dynamics of the modules annotated with various cellular responses (rKID:56 – inflammation [representing gene: *Icam1*], rKID:127 – TGFβ signalling [representing gene: *Tgfbr2*], rKID:3m – oxireductase and mitochondrial function [representing gene: *Mpc2*], rKID:140 – sugar transport and metabolism [representing gene: *Pfkp*]) (blue) compared with the dynamic of rKID:160- DNA damage (red). The representing genes indicate strongly modulated gene memberships with the functions associated to the module annotation. The stars represent the significant log2 fold change values (adj p-val < 0.05).

### Differential regenerative processes between CPT and OMPT

Renal injury and death of proximal tubular epithelial are counter balanced by regeneration programs^53^. Our dataset allowed us to determine the spatio-temporal activity of the programs that drive the tissue regeneration and the involvement of different nephron segments in these processes. The injured kidneys underwent regeneration processes starting from day 3 onward (Suppl. figure 6A). Since tissue regeneration implies replacement of dead cells by new cells, we first focused on module rKID:5m which reflects the activity of the cell cycle related genes (Figure 7Ai and Suppl. table 5). The cell cycle activity increased drastically in the OMPT peaking at day 5; only minor activation of module rKID:5m at later time points was observed in CPT. Activation of this module in OMPT was consistent with an increase of the number of Ki67-positive nuclei peaking in the outer-medulla region between day 3 and day 12 (Figure 7Bi and ii). These data indicate the local initiation of cell cycle programs mainly in OMPT following the DNA damage and cell injury.

**Figure 7.**
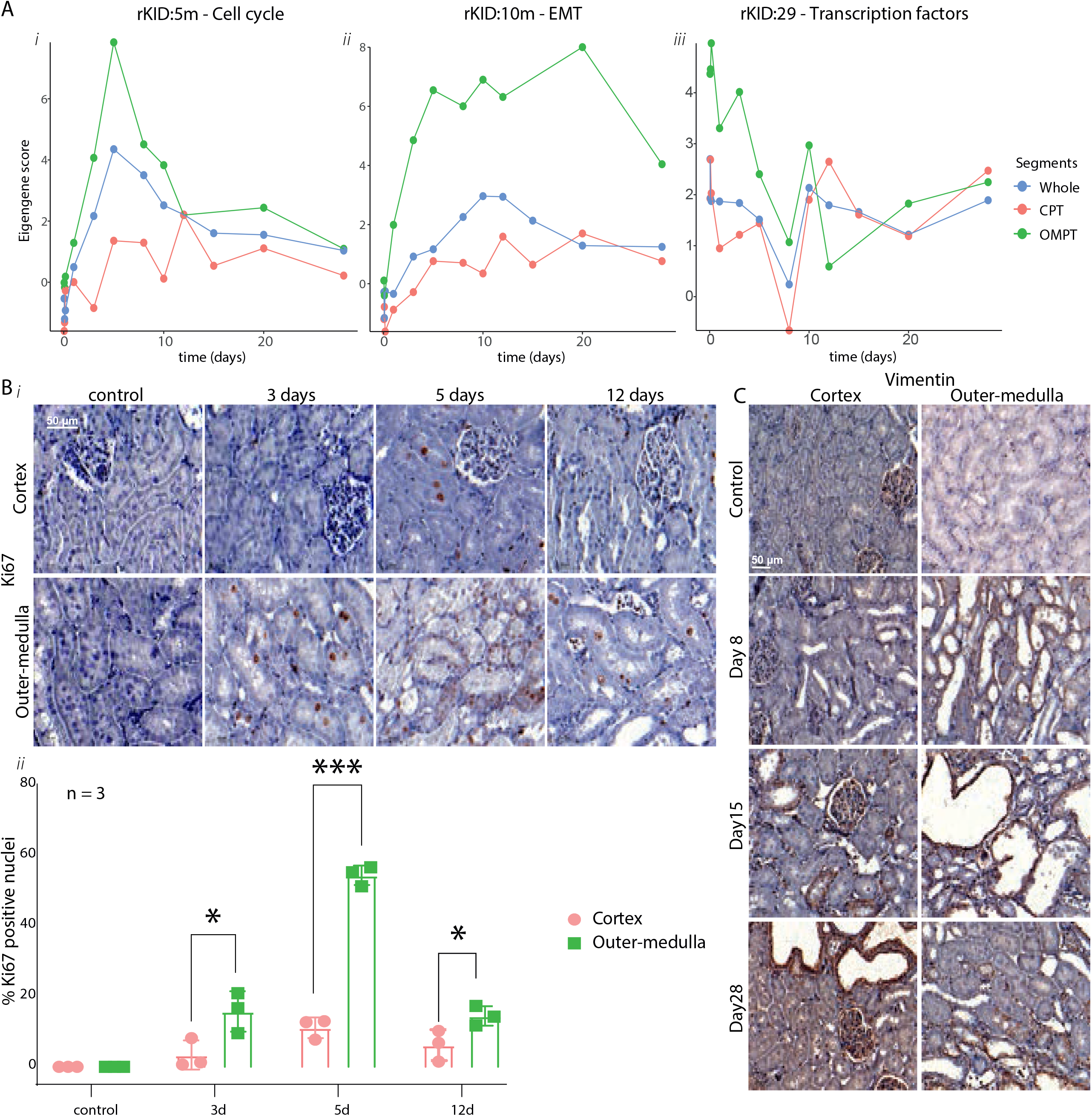
Temporal regenerative responses elicited in CPT and OMPT after cisplatin treatment. (A) Temporal dynamics of rKID:5m-Cell cycle (i), rKID:10m-epithelial-mesenchymal transition (ii) and, rKID:29-transcription factors (iii) in each sample type. (B) Nuclear staining of Ki67 in cortex (top) and outer-medulla (bottom) regions – 60x magnification (i). Quantification of Ki67 positive nuclei in the cortex (red) and outer-medulla area (green). The error bars indicate the standard deviation values. Each dot in the plot shows the value of each replicate. The stars indicate the p-values derived from independent student t-test indicate the significant comparison between cortex and outer-medulla regions (* p-val <0.05 – 0,01, ** p-val <0.01 – 0.001, p-val *** < 0.001). (D) Dynamics of in each sample type (ii). (C) Temporal dynamic plots of module in each sample type. (F) Immunohistochemistry staining of vimentin in the cortical (left) and outer-medulla (right) areas 40x magnification.

Epithelial-mesenchymal transition (EMT) and reorganization of the extracellular matrix are critical processes during renal injury responses and involved in wound healing, tissue regeneration, and organ fibrosis^54,55^. We examined module rKID:10m which has the strongest enrichment for extracellular matrix organization among all modules. The activation of rKID:10m occurred primarily in OMPT with a sustained high level between day 5 and 20 post cisplatin administration (Figure 7Aii). The expression of the mesenchymal marker *Vim*^56^ displayed a similar dynamic profile to the change in rKID:2m (its parent module) EGs (Figure 8A) with early and strong upregulation in OMPT and later contribution of CPT (Suppl. figure 7A). The transcriptional temporal dynamics of the local EMT program activation was supported by IHC of vimentin with strong involvement of outer-medulla regions and CPT that underwent injury (Figure 7C and Suppl. figure 7B and C). Both sustained activation of module rKID:5m and vimentin expression till 28 days, with regeneration/degeneration and fibrosis indicated that the kidneys were not yet fully restored till end of the study.

**Figure 8.**
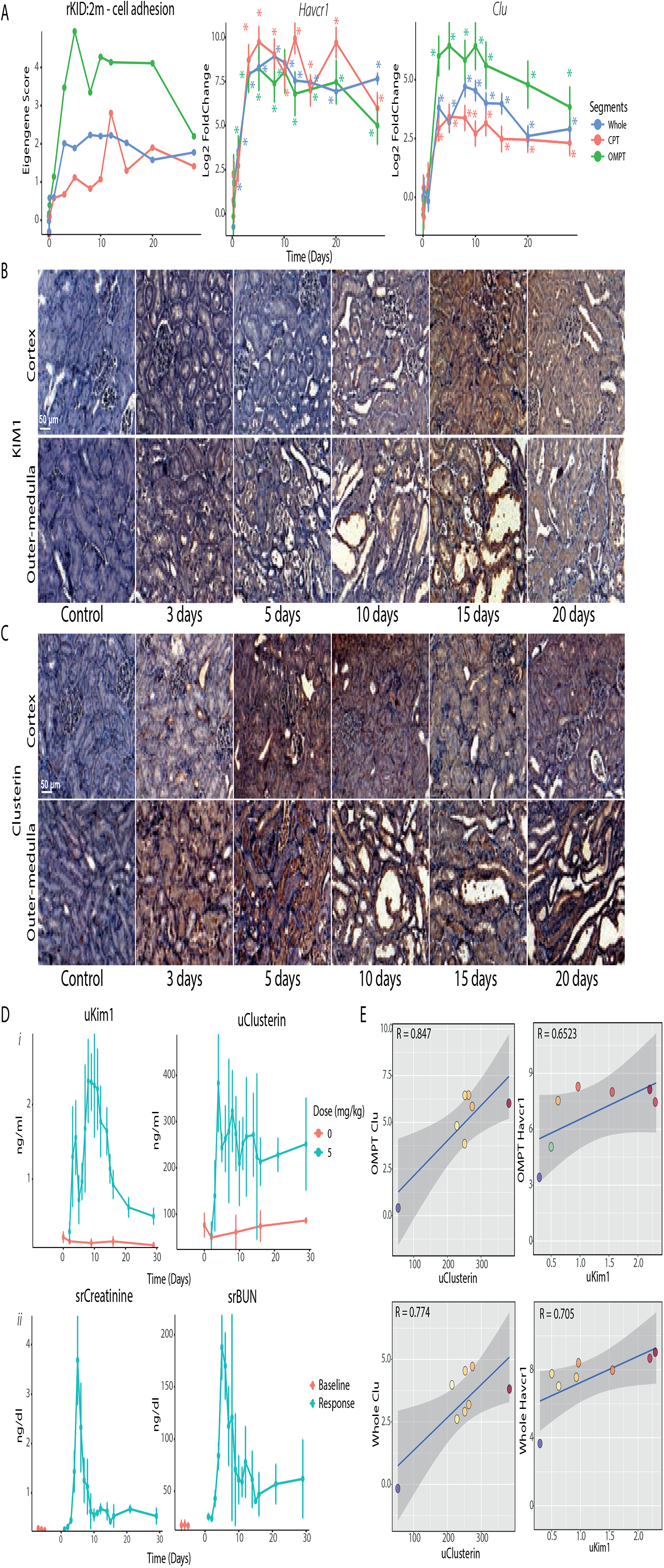
Temporal expression of kidney injury biomarkers upon cisplatin exposure. (A) Temporal dynamic plots of rKID:2m – Kidney injury biomarkers in the whole kidney (blue), CPT (red), and OMPT (green) – (i). Time dynamic plots of *Havcr1* (ii) and *Clu* (iii) two kidney biomarker genes in each sample type (error bars represent SEM) - right. The stars represent the significant log2 fold change values (adj p-val < 0.05). (B) Immunohistochemistry staining of KIM1 and in the cortical (upper) and outer-medulla (lower) region – 40x magnification. (C) Immunohistochemistry staining of clusterin and in the cortical (upper) and outer-medulla (lower) region – 40x magnification. (D) Temporal dynamics of urine KIM1 and clusterin (i) as mechanistic biomarkers and serum creatinine and BUN (ii) as functional biomarkers. Red plot represents the baseline measurement with error bars indicate standard deviation values. (E) Correlation plots between urine clusterin with *Clu* expression in OMPT (i-top), urine KIM1 with *Havcr1* expression in OMPT (ii-top), urine clusterin with *Clu* expression in the whole kidneys (i- bottom), urine KIM1 with *Havcr1* expression in the whole kidneys (ii-bottom).

The early cellular response activation in OMPT could be explained by the rapid and strong activation of rKID:29 in the OMPT (Figure 7Aiii). The top 3 gene memberships of this modules (*Btg2, Atf3*, and *Ier2*) also exhibited rapid upregulation in OMPT (Suppl. figure 6B). Our findings confirmed that the activation of the transcription factors linked to the early injury responses were likely associated with the strong and early modulation of cellular responses related to injury and regeneration.

### Urinary clusterin reflects sustained OMPT injury

The detailed mechanistic insight in the spatio-temporal activation of both initial damage responses as well as regeneration/degeneration programs allowed us to more accurately define their relationship with standard clinical chemistry and candidate novel kidney biomarkers. Module rKID:2m, which contained inducible kidney biomarker genes (see above), showed a strong activation in OMPT with rapid and sustained temporal activation compared to whole kidney and CPT (Figure 8A-left). Interestingly, the expression of biomarkers *Havcr1* (KIM1) and *Clu* (clusterin) (Figure 8A-middle and right) exhibited different regional specificity. The high upregulation of *Havcr1* was comparable in both CPT and OMPT, whereas the expression of *Clu* was more strongly increased in OMPT. We further verified the transcriptomics data with an immunostaining of the respective proteins KIM1 and clusterin. KIM1 protein expression showed indeed similarity in both the cortical and outer-medulla areas (Figure 8B), while clusterin staining was much stronger and occurred earlier (already at day 3) in the outer-medulla region (Figure 8C). The uKIM1 and clusterin (Figure 8Di) both showed a strong elevation from day 2 onward (Figure 8Di). Yet, uKIM1 peaked at day 10 and decreases rapidly, while uClusterin remained remarkably high until day 28 consistent with the kidney tissue immunostaining (Figure 8C – supplementary figure 8). The standard renal clinical chemistry markers srCreatinine and srBUN showed a later onset than uKIM1 and uClusterin and peaked at day 3; thereafter a rapid decline was observed (Figure 8Dii), regardless of sustained kidney pathology (Suppl. figure 6). In addition, uClusterin highly correlated with *Clu* expression in the injured OMPT and in whole kidney (Figure 8Ei), while uKIM1 showed lower correlation with *Havcr1* expression in OMPT (Figure 8Eii). These results demonstrate that the mechanistic biomarkers more accurately reflect the temporal molecular events and histopathology, with clusterin being the most representative for the severe spatial cisplatin-induced injury response in OMPT.

## Discussion

In the current study we performed a detailed and systematic analysis of the spatial-temporal dynamics of the cell signaling responses in different nephron segments upon cisplatin exposure. First, we established and validated a new protocol combining laser capture micro-dissection with targeted RNA-sequencing and subsequent gene co-expression network activity analysis. Our findings indicate that: 1) OMPT, and not CPT and GM, are the direct primary target of cisplatin with a sustained DNA damage response; 2) The DNA damage response in OMPT is parallel to the activation of inflammatory responses, cytoskeletal remodeling and glycolytic reprogramming, the latter associated with repression of oxidative mitochondrial function; 3) the regeneration programs are particular activated within the injured nephron segment; and, 4) urinary clusterin is most representative for the sustained OMPT injury. We have summarized these distinct spatial-specific responses of cisplatin nephrotoxicity in Figure 9.

**Figure 9.**
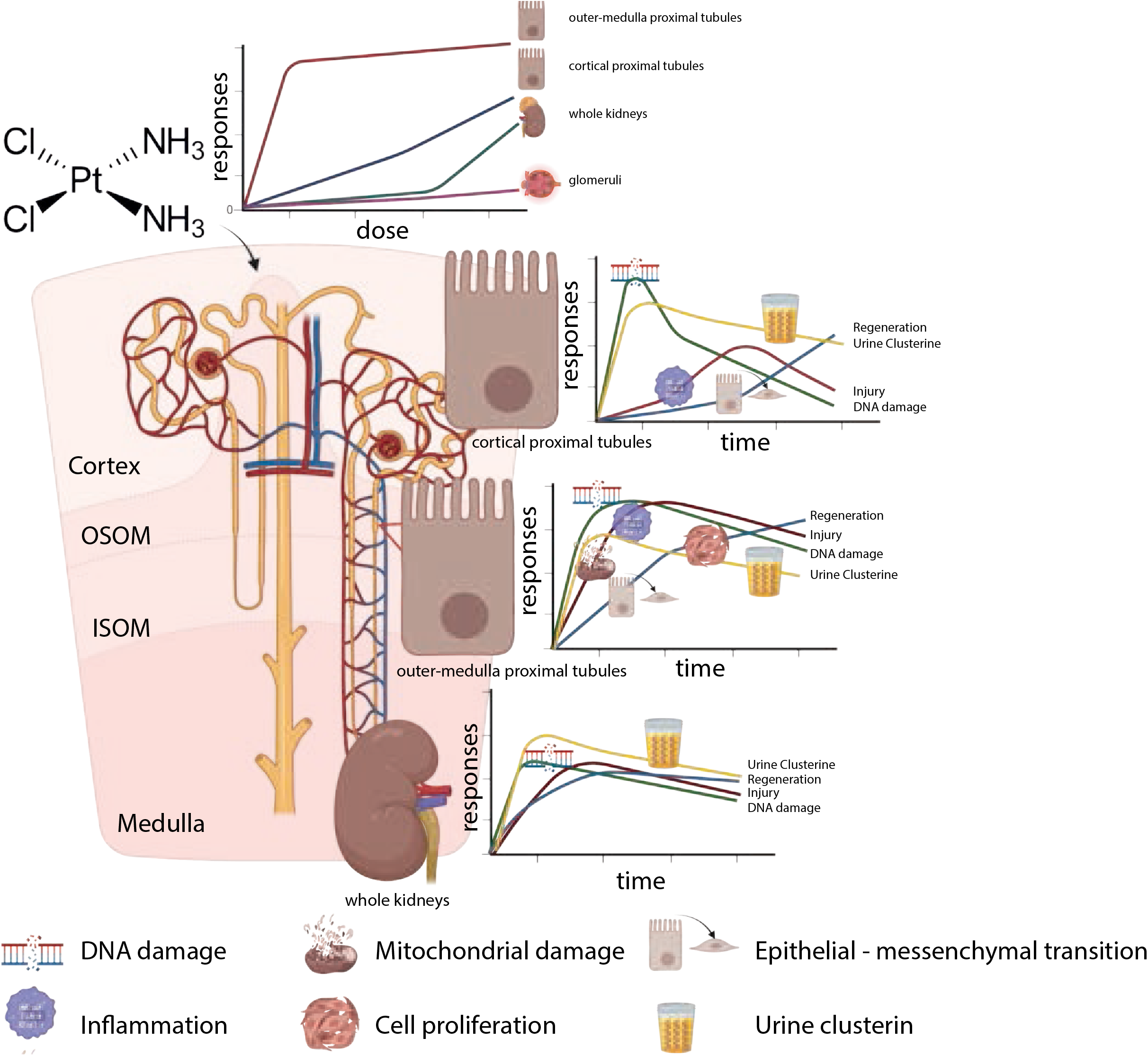
Schematic overview of cisplatin-induced key cellular responses elicited in the different segments of the rat kidneys. The difference in activation pattern of cellular responses is highlighted in whole kidney, CPT, and OMPT. This figure was created with BioRender.com.

Our LCM-TempO-Seq approach for spatial assessment of differential transcriptional responses in relation to kidney injury responses has not been applied previously. While we cannot fully exclude contamination of LCM samples with some neighboring cells, - e.g., CPT with distal tubule -, we have successfully demonstrated the strong region-specific enrichment of nephron segment-specific markers in the different LCM samples based on single cell RNA-sequencing data from mouse kidney^24^. The TempO-Seq outcomes provided sufficient read depth and gene coverage for integration with our Kidney TXG-MAPr tool for gene network analysis. The Kidney TXG-MAPr represents most renal patho- biological responses based on the transcriptional perturbations induced by various nephrotoxicants in association with diverse pathological conditions and spanning a time course of 3 hr to 29 days (manuscript in preparation). This platform provided clear qualification and quantification of gene network activities for the segment specific gene co-regulated network responses. Importantly, these responses were validated by independent immunohistochemistry of relevant representative markers of the key gene networks. We recognize that a major disadvantage of LCM is the laborious approach to derive detailed spatial transcriptomics information. Alternative methods could involve single cell sequencing or spatial transcriptomics technology^59^. Yet, besides the disadvantage of high costs for detailed studies as performed in our current study, overall, the transcript coverage and read depth of the latter two methods will most likely provide insufficient gene module coverage for integration with Kidney TXG-MAPr platform. Despite these disadvantages of LCM-TempO-Seq, we anticipate that our spatial transcriptome approach will increase the understanding of regional processes that are activated during (toxicant-induced) disease progression as well as efficacy of drug-induced disease regression.

OMPT was the most sensitive nephron region, with the most diverse and sustained cellular response activity after cisplatin treatment. The OMPT region mostly consists of the segment 3 of proximal tubule which is known to be severely injured by cisplatin^60^. This segment has high expression of OCT2 (*SLC22A2*) that transports cisplatin to the cells across the basolateral membrane^61^. Previous studies confirmed higher platinum concentration in the outer stripe of medulla compared to the cortex^62,63^. Given that the glomeruli are extracellularly exposed to cisplatin and show absent DNA damage responses further supports the requirement for active uptake of cisplatin in proximal tubular cells. Previously, *in vitro* primary cultured mouse proximal tubules of S3 segment were reported to be highly susceptible for cisplatin toxicity^60^. However, intriguingly, the initial activation of the DNA damage response in CPT was similar as in OMPT, but then later declined particularly in CPT (Figure 5B). This might suggest that initial total uptake in CPT is similar to in OMPT, but that the clearance of either cisplatin and/or DNA repair programs differ between CPT and OMPT. Lower expressions of Ctr2 (*SLC31A2*) which mediates the efflux of cisplatin through the apical site, may lead to higher platinum accumulation in the OMPT^64^. Our data indicates that the transcriptomics responses observed in whole kidney samples strongly underestimates the extent of cellular responses. Given that cisplatin only drives biological programs in a limited region of the kidney would naturally lead to the attenuation of these transcriptional responses in whole kidney perspectives. This directly highlights the power and necessity of the spatial analysis of transcriptional responses, since it would more precisely and sensitively uncover transcriptional programs that drive renal injury. Focusing solely on the responses of whole kidneys might easily overlook the actual responses in the affected regions.

Our data showed earlier and stronger upregulation of transcription factors initiating the responses of injury: ATF3, BTG2, and IER2 in OMPT compared to CPT. The activation of BTG2 was previously linked to cell death^32^. While ATF3 was found as a regenerative marker^34,57^. The upregulation of IER2 was reported to be involved in the differentiation processes^58^. This different activation pattern could delineate the different dynamics of following cellular responses e.g., DNA damage response, showing reversible activation in CPT, but sustained in OMPT. The stronger and sustained activation of the DNA damage response in the OMPTs was directly associated with the sustained activation of other processes such as inflammation, actin cytoskeletal reorganization, and a switch from mitochondrial activity to glycolysis. Multiple studies highlight the connection between overall severity of sustained DNA damage and progressive cellular responses^65–67^. The sustained DNA damage response in OMPT induced the continuous expression of *Cdkn1a*/P21 (Figure 5). Previously, the prolonged P21 expression as a consequence of the sustained DNA damage has also been found to hinder the proliferation activity impairing the regenerative process in the damaged kidneys^68,69^. Interestingly, P21-null mice also show reduced cisplatin nephrotoxicity^69^. Since in OMPT, the cell cycle repair and EMT programs are initiated by proximal tubule injury, blocking cell cycle progression due to sustained expression of *Cdkn1a* in the OMPT could be a critical factor that suppresses efficient regeneration of this region and contributes to a prolonged regeneration/degeneration phenotype. Altogether, our findings highlight the association between early and strong upregulation of the transcription factors with the activation of downstream cellular responses.

We discovered that urinary clusterin reflects more accurately the condition of OMPT than urinary KIM1. Previously, it has also been reported that the protein expression of clusterin is more sensitive than KIM1^70^. The onset of KIM1 expression was also seen in CPT suggesting that early activation of DNA damage response is sufficient for KIM1 activation but does not result in a drastic early onset of cell death. The measurement of the urinary clusterin could be a potential clinical biomarker to evaluate the overall progression of kidney injury in direct relation to sustained tissue injury. In our hands, this was related to specific injury to OMPT by cisplatin. Whether this also holds true for renal injury that is more specific for CPT needs further evaluation. While the increase of the serum creatinine and BUN demonstrated loss of renal function, these markers readily declined at later time points. This early decrease of renal filtration function could be caused by the physiological stress in glomeruli upon tubular damage that might contribute to the non-specific cellular response activation. This limitation of the serum biomarkers further highlights the necessity of measurement of mechanistic urinary biomarkers that more faithfully reflect the actual kidney injury at the moment of sampling. Moreover, the early transcriptional changes of these biomarker genes highly correlate with the urinary excretion of the proteins, especially for clusterin, suggesting the possibility of transcriptomic responses in predicting renal injury.

In summary, we have mapped the spatial-temporal transcriptional gene-network activation after cisplatin treatment using a novel LCM-TempO-Seq approach. Future studies using other nephrotoxic drugs that target other nephron segments would provide further insight in the variety of transcriptional programs in renal injury and regeneration and provide further direction to additional mechanism-based biomarkers. We anticipate that our methodology would allow a more refined understanding of biological responses in the kidney during drug development as well as in experimental models of renal disease progression and in the clinical setting.

## Author Contribution

L.S.W., S.J.K., P.T., C.F., M.E.C., D.C., K.S., S.L.E., J.L.S., and B.vd.W. designed the study; K.S., E.E.V., L.B., and T.R. carried out the *in vivo* experiments, performing biomarker analysis, and histological assessment; L.S.W and C.P. performed the laser capture microdissection and immunohistochemistry; L.S.W., S.A.B., and S.Z. conducted the ICP-MS methodology for platinum determination; L.S.W. and P.T. analyzed the transcriptomic data and biomarker results respectively; L.S,W., S.L.D., and B.vd.W. made the figures, drafted and revised the paper; all authors read and approved the final version of the manuscript. M.E.C, currently employed at Regeneron Pharmaceuticals, contributed to this article as an employee of AbbVie and the views expressed do not necessarily represent the views of Regeneron Pharmaceuticals Inc.

## Acknowledgement

This work was supported by the Innovative Medicines Initiative (IMI) 2 Joint TransQST project (grant agreement No 116030). The IMI Joint Undertaking receives support from the European Union’s Horizon 2020 research and innovation program and EFPIA. The authors would like to acknowledge Dr. Daniela Salvatori (Leiden University Medical Centrum) for the intellectual contribution during the development and optimization of laser capture microdissection methodology for the isolation of the nephron segments.

## Conflict of Interest

MC, currently employed at Regeneron Pharmaceuticals Inc, contributed to this article as an employee of Abbvie and the views expressed do not necessarily represent the views of Regeneron Pharmaceuticals Inc.

## Figure Legends

**Supplementary figure 1. Quality control output of the cisplatin dose-response *in vivo* experiment**.

(A) A bar plot showing different sequencing output of the frozen kidney section and FFPE kidney section based on the %mapped. The color of the bars indicate the preservation types (blue : cryopreservation and orange : FFPE). The error bar indicate standard deviation. (B) A principle component analysis (PCA) plot showing the two clusters of the samples with percentage (%) mapped higher than 10% (red) and lower than 10% (blue) (i). The sample exclusion plots based on the %mapped, with threshold 10% (ii). Every dot represents one sample and the color of the dots indicates the % mapped of each sample : higher than 10% (red) and lower than 10% (blue). (B) The plots show the Pearson correlation values of each sample from the particular condition (x-axis) with the mean from all the (biological) replicates in the same condition (i). Distribution plots of normalized count of the probes in every sample condition (each plot is derived from the normalized count value from all the sample with the same condition – biological replicates) (ii).

**Supplementary figure 2. Quality control output of the cisplatin temporal response *in vivo* experiment**. (A) A principal component analysis (PCA) plot showing the two distinctive clusters of the samples with percentage mapped higher than 10% (red) and lower than 10% (blue); The size of the dots represents the administered dose; The shape of every dot represents the sample type (i). The sample exclusion plots based on the %mapped with threshold 10%. Every dot represents the %mapped of each sample: higher than 10% (red) and lower than 10% (blue) (ii) (B) The correlation plots showing the pearson correlation value of each sample from the particular condition (x-axis) with the mean from all the (biological) replicates in the same condition. (i). Distribution plots of normalized count of the mapped probed in every sample condition (each plot consists of the normalized count value from all the sample with the same condition – biological replicates) (ii).

**Supplementary figure 3. Post normalization quality control output of the cisplatin temporal response *in vivo* experiment**. (A) The volcano plots of log2FC vs –log10 adjusted p-value of control conditions compared to the control condition at 672 hours (28 days). The red dots indicate the probes that are differentially expressed with the threshold adjusted p-value < 0.05 and log 2-fold change > [0.5]. (B) The module coverage plot indicating the number of overlapping genes from the BioSpyder Rat Whole Genome platform to the genes from the Kidney TXG-MAPr. The color legend indicates the coverage category with red > 75% coverage, blue 50% - 75% coverage, and green < 50% coverage. Each bar plot of the plot represents the coverage value of each module with total number of plots equal to 399 indicating 399 modules (x-axis).

**Supplementary figure 4. Kidney toxicogenomics map and histopathology findings after cisplatin treatment**. (A) The toxicogenomic map whole kidneys exposed to various dose of cisplatin ranging from 1-7.5 mg/kg) at 72 hours. Each circle on the map represents one module (highlighted: rKID:160- DNA damage module, detailed module membership overview of the network is displayed on the right). The color and the size of the circle indicate the modulation magnitude of the module responses (red: activation, blue: deactivation). (B) The histopathological readout of the kidney tissue focusing on the proximal tubules. The readout is split into 3 categories: degeneration/regeneration, karyomegali and tubular epithelial cells, and necrosis. Each category is measured by the degrees of severity (lower to higher). The intensity of the red color indicates the readout score in each category and the degree of severity assessed in every administered cisplatin dose.

**Supplementary figure 5. Specific kidney responses after cisplatin treatment**. (A) The expression of *Mgmt* (left) and *Nudt11* (right) upon cisplatin exposures at 72 hours. The color of the plots represents the expression of the genes in every segment of the nephrons. (B) KIM1 immunostaining in the cortex and outer-medulla region of the kidneys exposed to multiple doses of cisplatin – 40x magnification. (C) The dose-response plots of urine biomarkers (albumin, KIM1, and NGAL) and serum biomarkers (creatinine and BUN) in every time point. The colors of the plot indicate the collection time in day.

**Supplementary figure 6. Proximal tubule histopathology (A) and the upregulation of the top 3 highest module correlation gene memberships of rKID:29 (B)**. (A) The readout is split in 4 categories: necrosis, degeneration/regeneration, degeneration, and fibrosis. Each category is measured by degrees of severity (lower to higher). The intensity of the red color indicates the readout score in each category and the degree of severity assessed in every kidney collection time. (B) The time-response plots of 3 highest correlated gene members of rKID:29 in each sample type. The color of the plot shows the expression of gene memberships (error bars represent SEM). The stars represent the significant log2 fold change values (adj p-val < 0.05).

**Supplementary figure 7. Vimentin expression and immunohistochemistry of whole kidney, CPT and OMPT after cisplatin treatment**. (A) Expression of *Vim* as the mesenchymal marker in whole kidney (blue), CPT (red), and OMPT (green) (error bars represent SEM). The stars represent the significant log2 fold change values (adj p-val < 0.05). (B) The immunostaining of vimentin on the cortex (top) and outer-medulla region (bottom) at 5 days (left) - 40x magnification. (C) Vimentin expression in the CPTs pointed with the black arrows are found at the close proximity to the medullary ray (the area within red lines) (right) - 20x magnification.

**Supplementary figure 8. Additional clusterin staining**. The immunostaining of clusterin (right) at 28 days on the cortex (top) and outer-medulla region (bottom) - 40x magnification.

**Supplementary table 1. Log2 fold change values of dose-response study**

**Supplementary table 2. Log2 fold change values of temporal dynamics study**. The outcomes of the OMPT at 15 days were excluded due to low %mapped of the samples.

**Supplementary table 3. Module scores of dose-response study**

**Supplementary table 4. Module scores of temporal dynamics study**. The outcomes of the OMPT at 15 days were excluded due to low %mapped of the samples.

**Supplementary table 5. Module memberships of the modules linked to the mechanisms of cisplatin induced DIKI (modules which are investigated in this study)**

**Supplementary table 6. The information of the selected modules that are evaluated in the manuscript**. The modules were selected based on the annotation linked the known mechanisms of cisplatin induced kidney injury. The dose-response and time-response pattern of the modules were complementarily evaluated to ensure that the responses of the modules potentially represented the biology of the affected tissue.

**Supplementary table 7. Nephron markers obtained from the previous study**^24^

**Supplementary table 8. The top 10 modulated gene networks in GM upon the exposure of 7.5 mg/kg of cisplatin**.

## Notes

### Competing Interest Statement

The authors have declared no competing interest.

