## supplementary table for "Spatio-temporal transcriptomic analysis reveals distinct nephrotoxicity, DNA damage and regeneration response after cisplatin": SupplementaryTable6_Selected_Module_identity.docx

| Module | Nr_genes | Hub_gene | EntrezID | cor_EG | Key_annotation | Annotation_lvl1 | Annotation_lvl2 | Annotation_lvl3 |
| --- | --- | --- | --- | --- | --- | --- | --- | --- |
| rKID:2m | 1123 | Tes | 500040 | 0.915 | Cytoskeleton; cell adhesion / injury biomarkers | Cytoskeleton | Cell adhesion | Actin filament organization |
| rKID:3m | 609 | Akr7a2 | 171445 | 0.914 | Metabolism | Metabolism | Small molecule metabolic process | Organic acid metabolism |
| rKID:5m | 318 | Mad2l1 | 297176 | 0.927 | Cell cycle; S/M phase | Cell cycle | Chromosome | Mitotic cell cycle |
| rKID:10m | 118 | Fbn1 | 83727 | 0.925 | Extracellular matrix (ECM) | Extracellular matrix | Extracellular matrix organization | Collagen formation |
| rKID:15 | 77 | Cyp2c13 | 171521 | 0.931 | Immune response; complement and coagulation | Immune response | Complement and coagulation | Enzyme inhibitor activity |
| rKID:26m | 64 | RGD1563941 | 500993 | 0.835 | Mitochondrion; translation | Mitochondrion | Mitochondrial translation |  |
| rKID:29 | 37 | Btg2 | 29619 | 0.93 | Transcription | Transcription | Transcription factors |  |
| rKID:56 | 24 | Icam1 | 25464 | 0.906 | Immune response; TNF/NFkB signaling | Immune response | TNF signaling | Response to bacterium |
| rKID:127 | 15 | Elk3 | 362871 | 0.885 | Cell migration | Cell migration | Epithelial cell migration |  |
| rKID:138 | 14 | Cd3g | 300678 | 0.904 | Immune response; T-cell receptor | Immune response | T-cell receptor signaling | Lymphocyte activation |
| rKID:140 | 14 | St14 | 114093 | 0.84 | Transport | Transport | SLC-mediated transmembrane transport |  |
| rKID:160 | 13 | Mdm2 | 314856 | 0.884 | Stress response; DNA damage / p53 | Stress response | DNA damage response | p53 pathway |
| rKID:280 | 9 | Ptpn12 | 117255 | 0.801 |  | NA |  |  |
| rKID:298 | 8 | Glrx3 | 58815 | 0.857 | Stress response; DNA damage / p53 | Stress response | DNA damage response | p53 pathway |

**Supplementary table 6. The information of the selected modules that are evaluated in the manuscript**. The modules were selected based on the annotation linked the known mechanisms of cisplatin induced kidney injury. The dose-response and time-response pattern of the modules were complementarily evaluated to ensure that the responses of the modules potentially represented the biology of the affected tissue.
