## supplementary figure for "Spatio-temporal transcriptomic analysis reveals distinct nephrotoxicity, DNA damage and regeneration response after cisplatin"

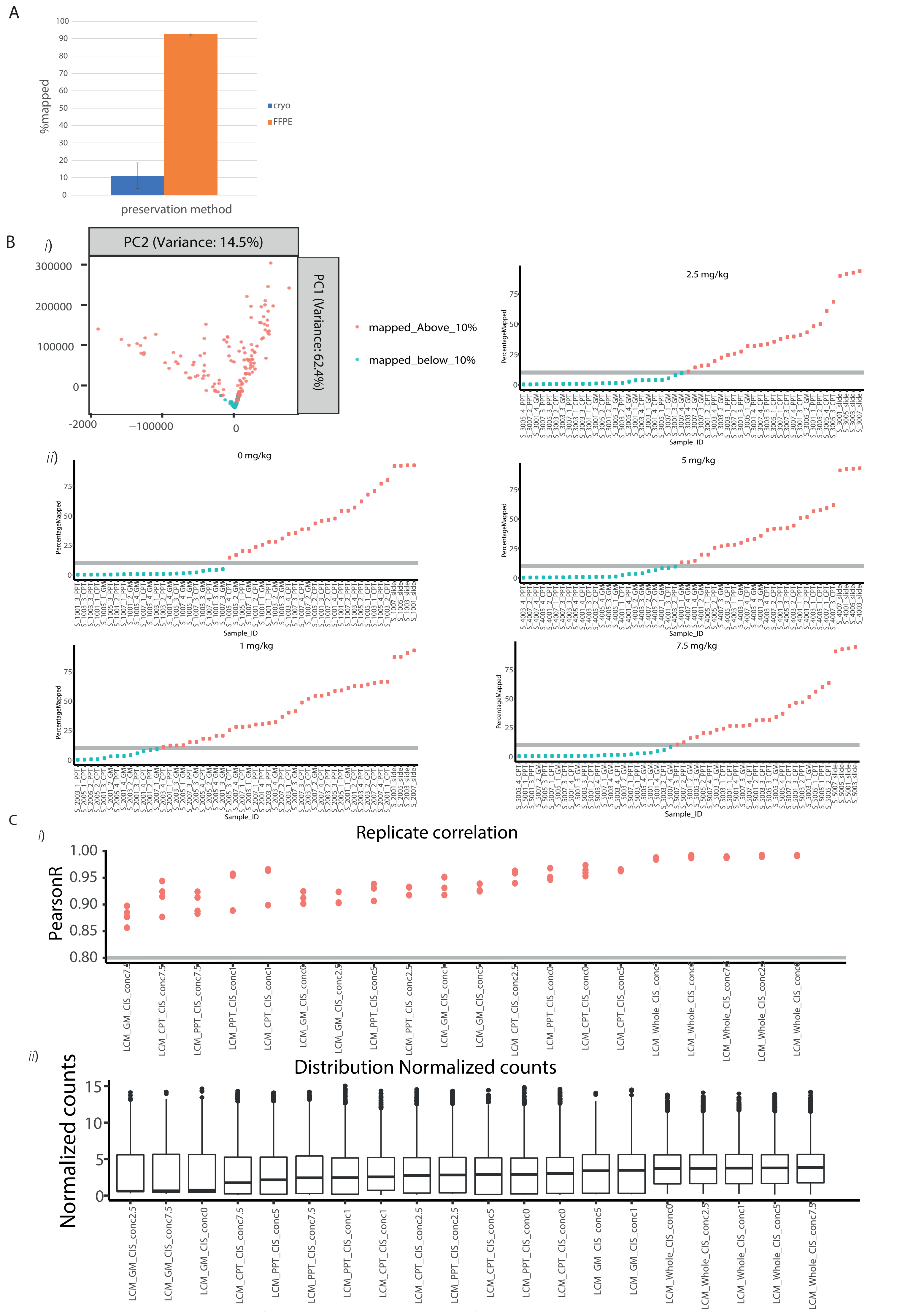


Supplementary figure 1. Quality control output of the cisplatin dose-response in vivo experiment

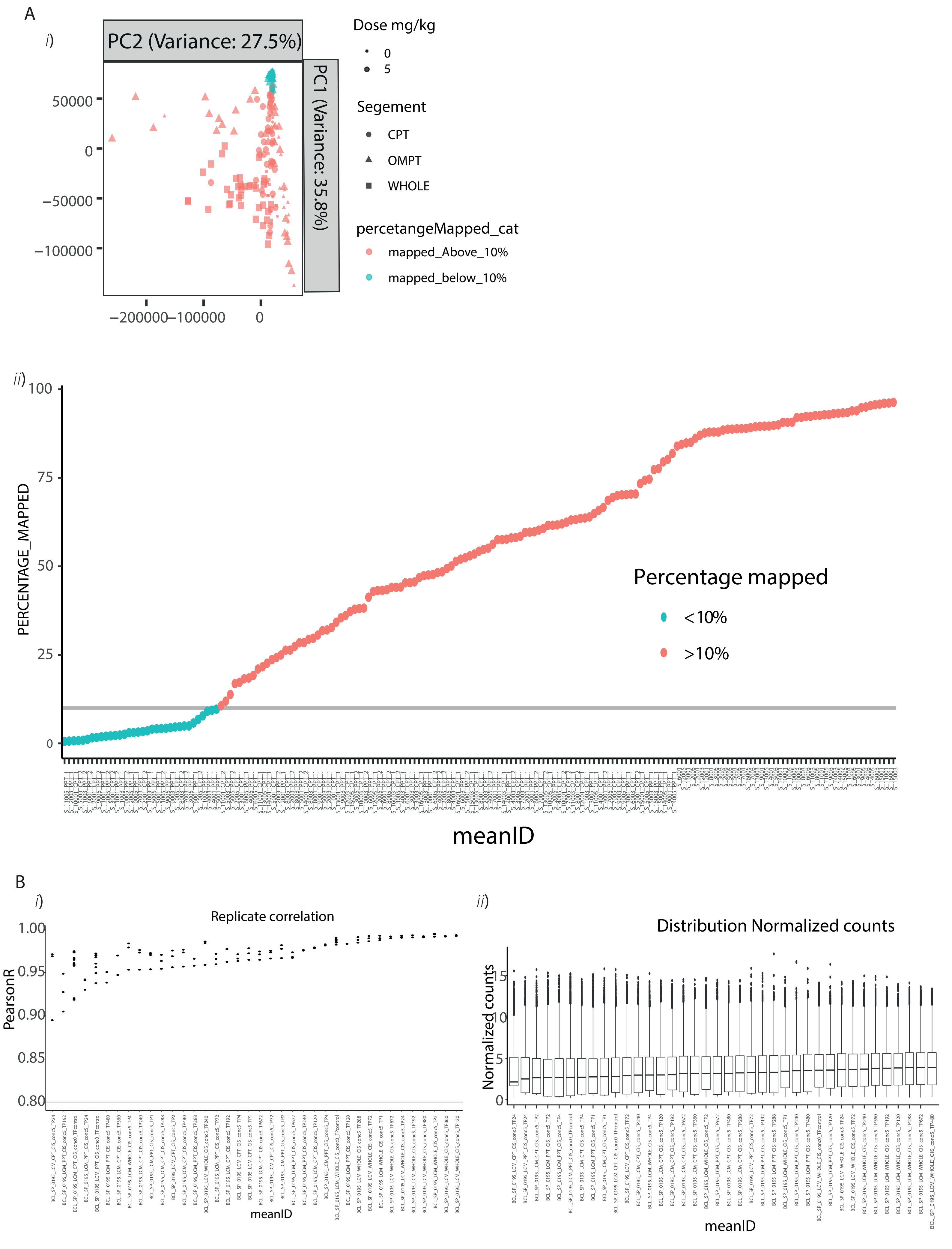


Supplementary figure 2. Quality control output of the cisplatin temporal response in vivo experiment

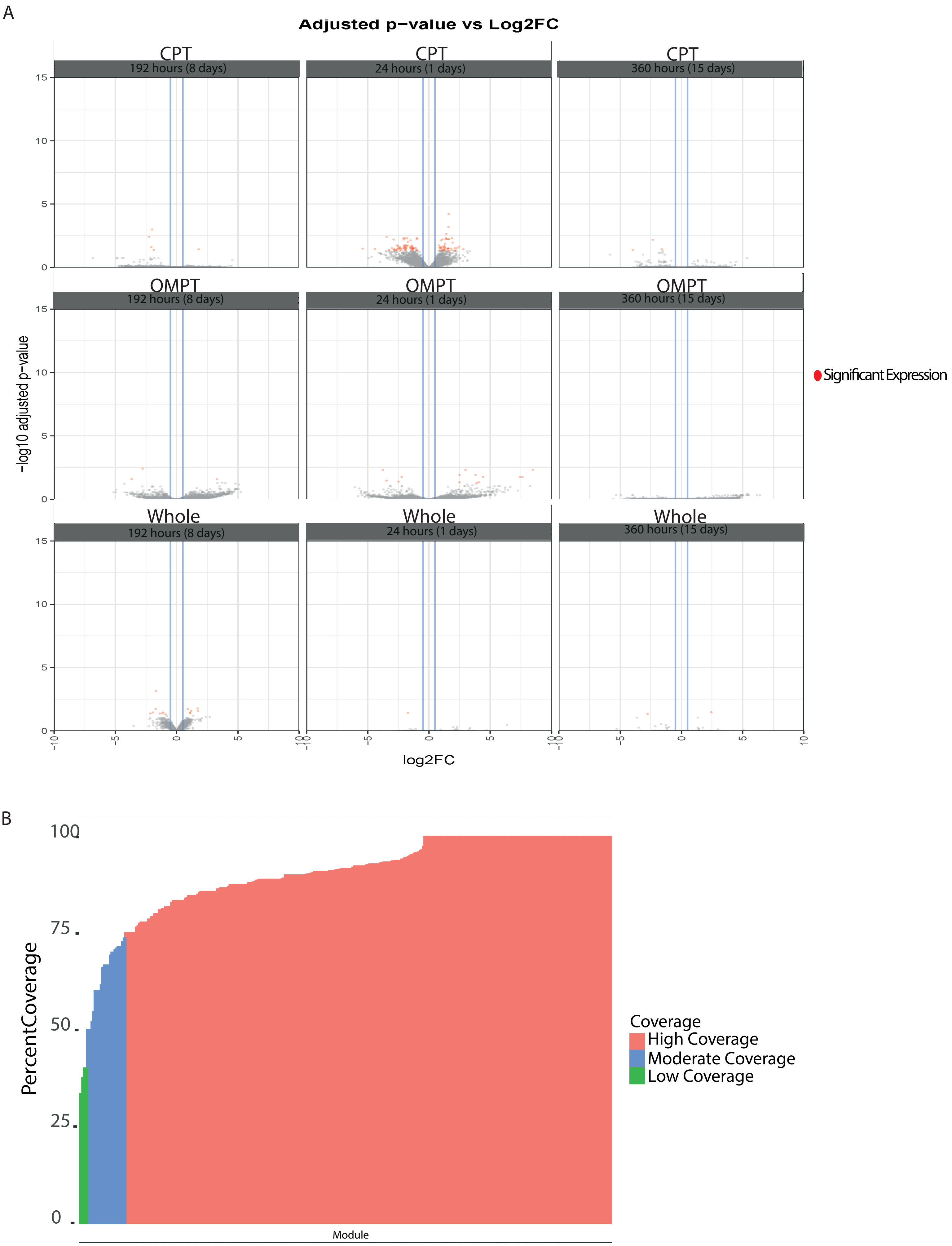


Supplementary figure 3. Post normalization quality control output of the time-dynamics in vivo experiment

A

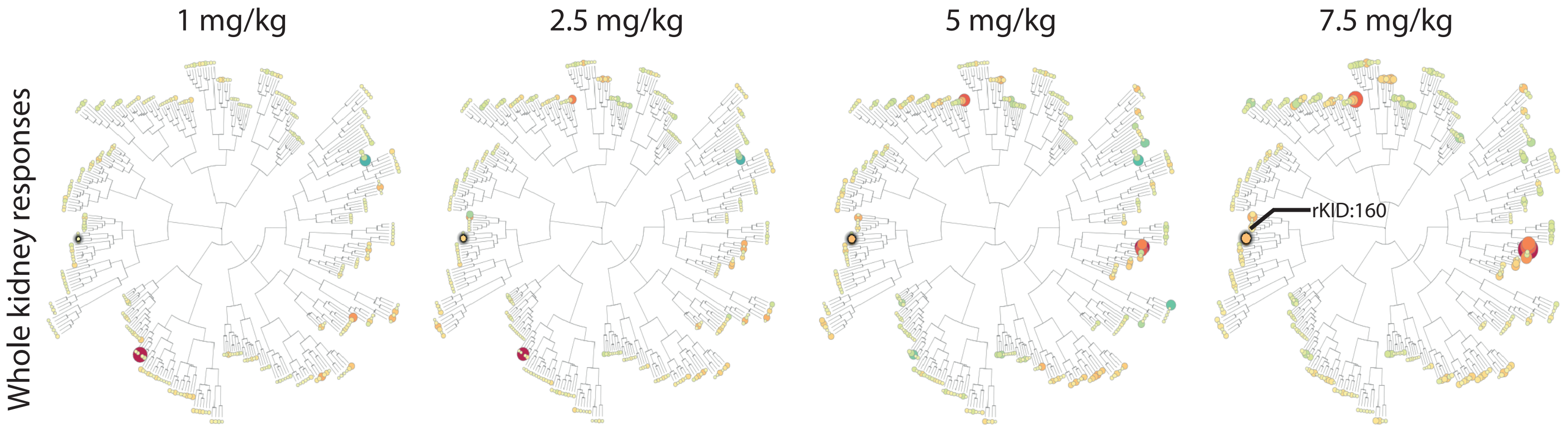


B

| Responses | Degree of Responses | 0 mg/kg | 1 mg/kg | 2.5 mg/kg | 5mg/kg | 7.5 mg/kg |
|---|---|---|---|---|---|---|
| Degeneration/Regeneration | minimal, focal, proximal tubule, unilateral | 0 | 1 | 0 | 0 | 0 |
| | minimal, multi-focal, proximal tubule, unilateral | 0 | 1 | 0 | 0 | 0 |
| | mild, multi-focal, proximal tubule, unilateral | 0 | 0 | 4 | 4 | 3 |
| | marked, multi-focal, proximal tubule, unilateral | 0 | 0 | 0 | 0 | 1 |
| Karyomegaly, tubular epithelial cell | minimal, focal, proximal tubule, unilateral | 0 | 2 | 0 | 0 | 0 |
| | mild, multi-focal, proximal tubule, unilateral | 0 | 1 | 1 | 1 | 4 |
| | moderate, multi-focal, proximal tubule, unilateral | 0 | 0 | 3 | 3 | 0 |
| Necrosis | minimal, multi-focal, proximal tubule, unilateral | 0 | 0 | 4 | 4 | 0 |
| | mild, multi-focal, proximal tubule, unilateral | 0 | 0 | 0 | 0 | 3 |
| | moderate, multi-focal, proximal tubule, unilateral | 0 | 0 | 0 | 0 | 1 |

Supplementary figure 4. Kidney toxicogenomics map and histopathology findings after cisplatin treatment

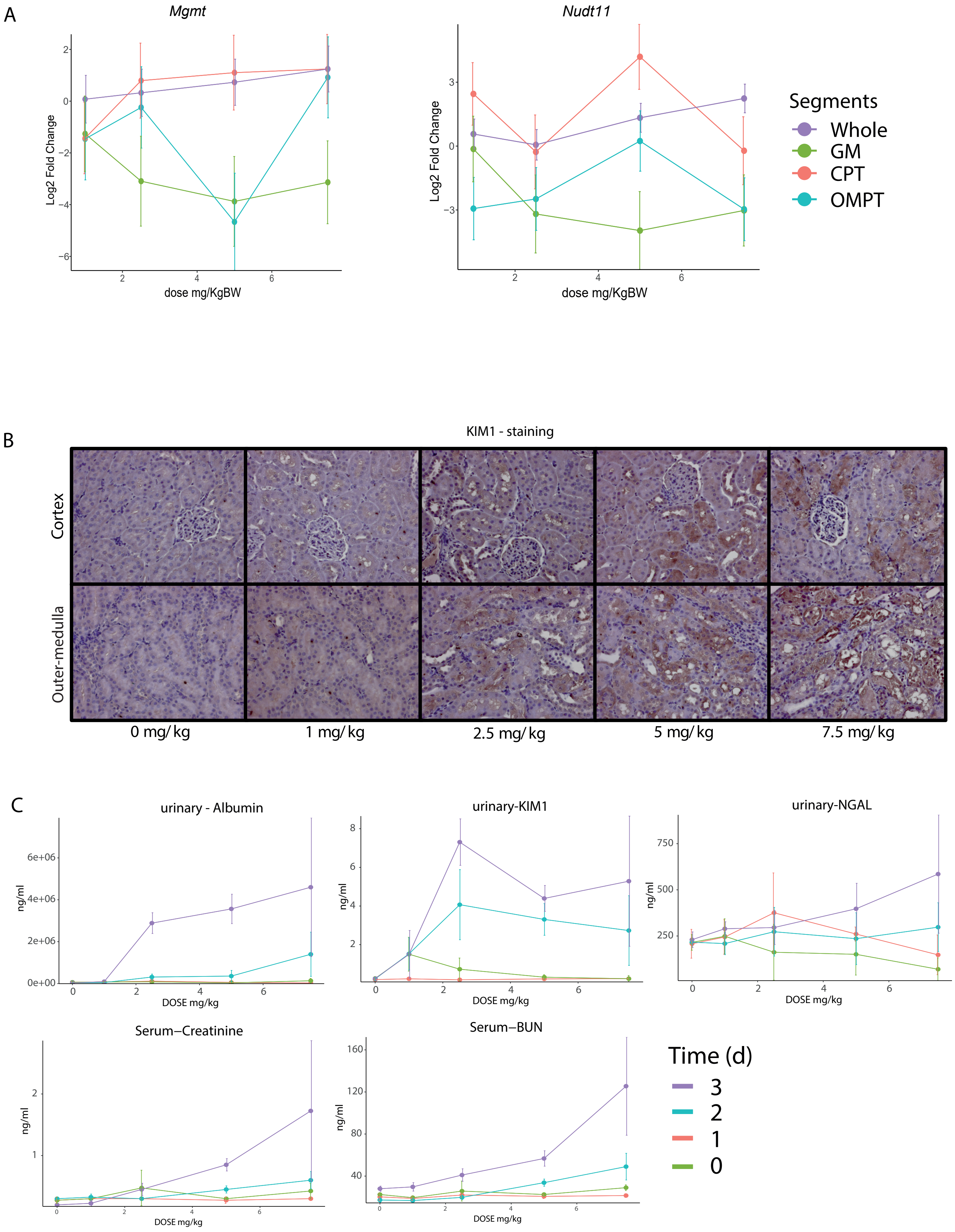


Supplementary figure 5. Specific kidney responses after cisplatin treatment

A

| Renal tubular findings | Grade / Day | 0 | 1 | 3 | 5 | 8 | 10 | 12 | 15 | 20 | 28 |
|---|---|---|---|---|---|---|---|---|---|---|---|
| Necrosis | minimal | 0 | 1 | 0 | 0 | 3 | 3 | 3 | 3 | 3 | 0 |
| | mild | 0 | 0 | 3 | 0 | 0 | 0 | 0 | 0 | 0 | 0 |
| | moderate | 0 | 0 | 0 | 3 | 0 | 0 | 0 | 0 | 0 | 0 |
| | marked | 0 | 0 | 0 | 0 | 0 | 0 | 0 | 0 | 0 | 0 |
| Degeneration/ Regeneration | minimal | 0 | 0 | 0 | 0 | 0 | 0 | 0 | 0 | 0 | 0 |
| | mild | 0 | 0 | 2 | 0 | 0 | 0 | 0 | 0 | 0 | 0 |
| | moderate | 0 | 0 | 1 | 3 | 0 | 0 | 1 | 2 | 1 | 2 |
| | marked | 0 | 0 | 0 | 0 | 3 | 3 | 2 | 1 | 2 | 1 |
| Degeneration | minimal | 0 | 0 | 0 | 0 | 1 | 1 | 0 | 0 | 1 | 1 |
| | mild | 0 | 0 | 0 | 0 | 2 | 1 | 1 | 1 | 2 | 0 |
| | moderate | 0 | 0 | 0 | 0 | 0 | 0 | 2 | 1 | 0 | 2 |
| | marked | 0 | 0 | 0 | 0 | 0 | 0 | 0 | 0 | 0 | 0 |
| Fibrosis | minimal | 0 | 0 | 0 | 0 | 1 | 0 | 3 | 3 | 1 | |
| | mild | 0 | 0 | 0 | 0 | 0 | 0 | 0 | 0 | 2 | 1 |
| | moderate | 0 | 0 | 0 | 0 | 0 | 0 | 0 | 0 | 0 | 2 |
| | marked | 0 | 0 | 0 | 0 | 0 | 0 | 0 | 0 | 0 | 0 |

B

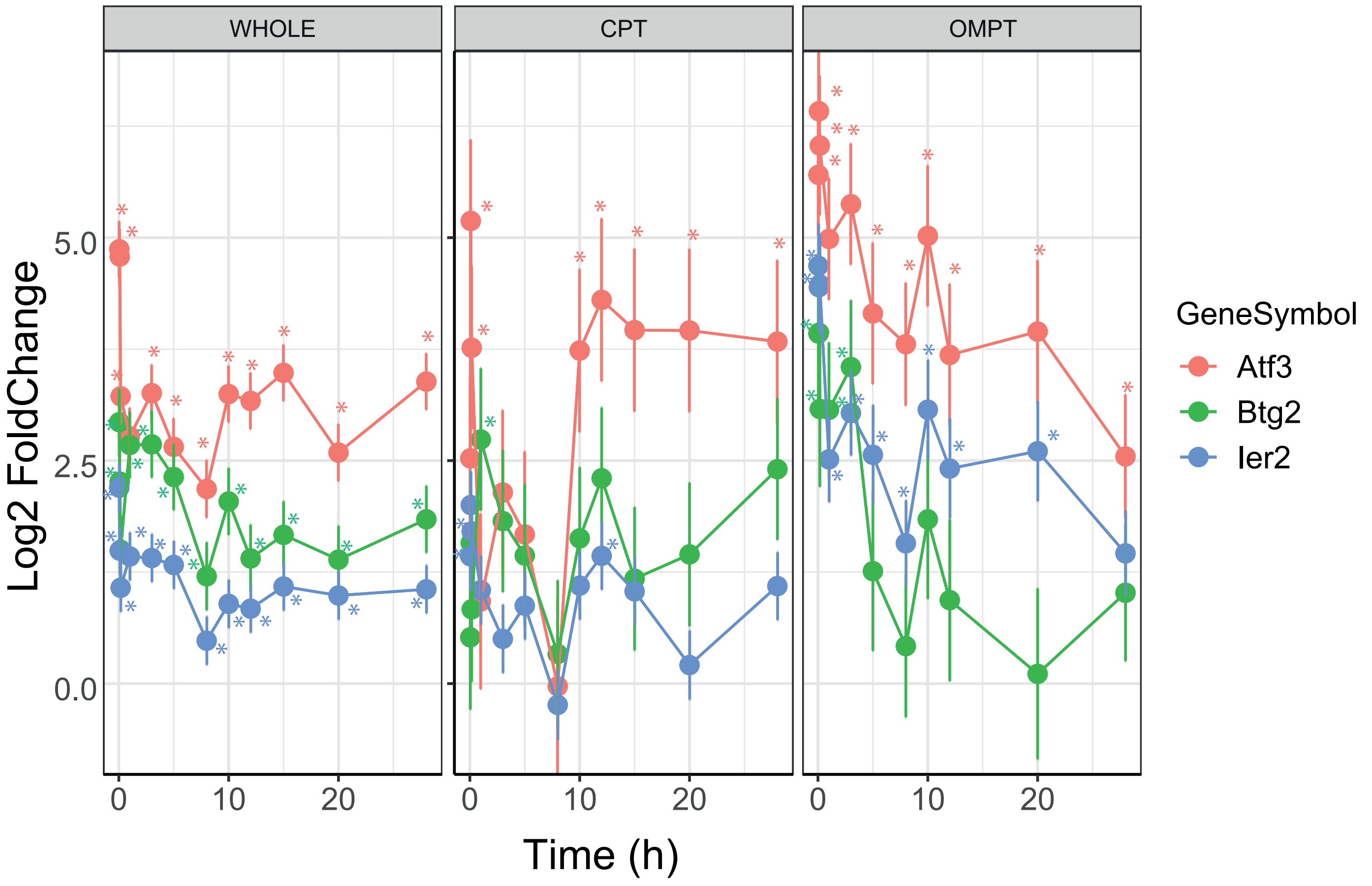


Supplementary figure 6. Proximal tubule histopathology (A) and the upregulation of the top 3 highest module correlation gene memberships of rKID:29 (B).

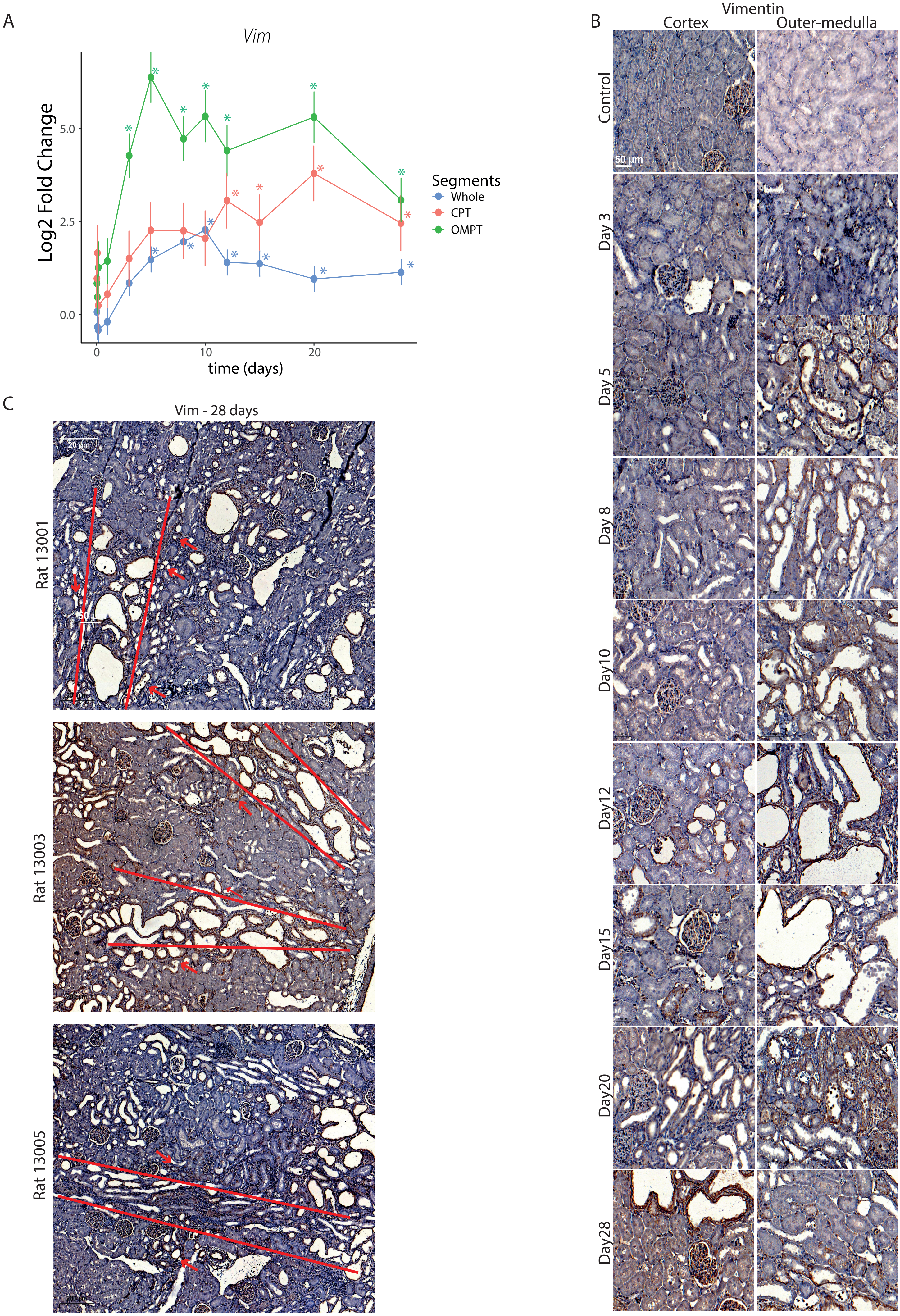


Supplementary figure 7. Vimentin expression and immunohistochemistry of whole kidney, CPT and OMPT after cisplatin treatment

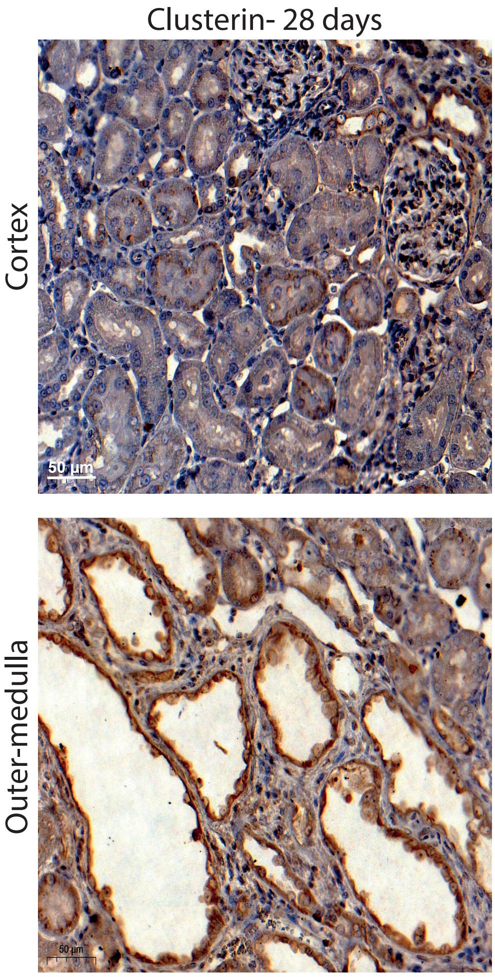


Supplementary figure 8. Additional clusterin staining
